# Liver Zonation Disruption Fuels Hepatocellular Carcinoma in Chronic Liver Disease

**DOI:** 10.64898/2025.12.06.692782

**Authors:** Sue Bin Yang, Federico Di Tullio, Lida Yang, Bruno Cogliati, Desiree Schnidrig, Andrej Benjak, Shamsa Roshan, Joana I. Almeida, Lilian Li, Sai Ma, Jan S. Tchorz, Charlotte K Y Ng, Tianliang Sun

**Affiliations:** Department of Stem Cell Biology and Regenerative Medicine, Icahn School of Medicine at Mount Sinai, New York, NY 10029, USA; Division of Liver Diseases, Icahn School of Medicine at Mount Sinai, New York, NY 10029, USA; Black Family Stem Cell Institute, Icahn School of Medicine at Mount Sinai, New York, NY 10029, USA; Mount Sinai Tisch Caner Certer, Icahn School o f Medicine at Mount Sinai, New York, NY 10029, USA; Department of Genetics and Genomic Sciences, Icahn School of Medicine at Mount Sinai, New York, NY 10029, USA; Department of BioMedical Research (DBMR), University of Bern, Bern, Switzerland; Institute of Physiology I, University of Tübingen, Tübingen 72076, Germany; Department of Biomedical Sciences, Humanitas University, Pieve Emanuele, MI 20072 Italy; IRCCS Humanitas Research Hospital, Rozzano, MI 20089, Italy

**Keywords:** liver metabolic zonation, regeneration, ZNRF3, RNF43, hepatocellular carcinoma, β-Catenin, hepatocyte identity

## Abstract

Hepatocellular carcinoma (HCC) arises almost exclusively in chronic liver disease (CLD), yet the classical etiological drivers of injury insufficiently explain why only a subset of patients progress to cancer. Here, we identify disruption of liver metabolic zonation, specifically, aberrant expansion of β-catenin activity from pericentral to periportal territories as a previously unrecognized tumorigenic risk state that emerges across etiologically diverse CLDs. Using spatial transcriptomics and immunohistochemistry from murine liver disease followed by human sample validation, we demonstrate that MASLD/MASH, alcohol-associated hepatitis, viral hepatitis, and immune-mediated cholangiopathies share a striking periportal induction of pericentral β-catenin target programs, indicating a conserved zonation disturbance independent of disease etiology.

Since β-catenin expansion occurs alongside inflammation, fibrosis, and metabolic dysfunction in human CLD, its direct oncogenic contribution remained unclear. To functionally isolate the tumorigenic consequence of zonation disruption itself from these confounding disease processes, we employed hepatocyte-specific deletion of ZNRF3 and RNF43—negative regulators that physiologically restrict β-catenin to the pericentral zone. Lineage tracing demonstrates that ZNRF3/RNF43 deletion drives selective periportal hepatocyte proliferation, zonal reprogramming, and tumor initiation in a β-catenin-dependent manner, establishing zonation disruption as a direct mechanistic driver of carcinogenesis. The resulting tumors exhibit a distinct metabolic and immunologic phenotype, including heightened mitochondrial respiration, preserved periportal identity, and T-cell competence, and correspond to a molecularly defined subset comprising 5-10% of human HCCs.

Together, these findings reveal β-catenin zonation expansion as a conserved and previously unrecognized HCC risk factor that mechanistically links chronic liver injury to malignant transformation. They further establish ZNRF3/RNF43 deletion as a tractable model of zonation-driven hepatocarcinogenesis and identify a distinct human HCC subtype with unique therapeutic vulnerabilities, opening new avenues for mechanism-based risk stratification, early detection and preventive therapeutic strategies in patients with chronic liver disease.

## Introduction

Hepatocellular carcinoma (HCC) is a leading global cause of cancer-related death worldwide and most often arises in the setting of chronic liver disease (CLD)^1–7^. Classical etiological drivers, including metabolic dysfunction–associated steatotic liver disease (MASLD), metabolic dysfunction–associated steatohepatitis (MASH), chronic viral hepatitis, alcohol-associated liver disease, and immune-mediated cholangiopathies, create inflammatory and fibrotic environments that increase susceptibility to malignant transformation^8–16^. However, these factors alone do not fully explain why only a fraction of patients with comparable degrees of liver injury or cirrhosis ultimately progress to cancer^1–3,9,17^. This discrepancy strongly suggests the existence of additional, poorly understood biological risk states that modulate tumor initiation during chronic liver injury.

One such candidate is liver metabolic zonation, the spatial partitioning of hepatocyte functions along the porto-central axis. Zonation ensures efficient metabolic homeostasis^18–20^ and is maintained by opposing gradients of WNT/β-catenin signaling, high in pericentral hepatocytes and actively suppressed in periportal hepatocytes^21–26^. Disruption of this architecture has been reported in MASLD/MASH, where periportal hepatocytes aberrantly acquire pericentral metabolic programs^27–29^. However, whether zonation dysfunction itself contributes to tumor susceptibility remained unclear.

Recent studies provided compelling evidence that the hepatocyte’s original zonal identity shapes its tumorigenic potential, positioning zonation as a fundamental determinant in liver cancer biology^30,31^. These finding rise a key unanswered question: does loss of proper zonation, rather than the specific etiology of injury, create a tumor-permissive state during chronic liver injury? Moreover, whether such zonation defects are conserved across etiologically diverse CLDs has not been systematically explored.

ZNRF3 and RNF43, two transmembrane E3 ubiquitin ligases, are critical gatekeepers that restrict WNT/β-catenin signaling to pericentral niche by promoting Frizzled receptors turnover^24,32,33^. Their loss results in ectopic expansion of β-catenin activity into periportal territories and reprograms periportal hepatocytes toward pericentral-like states^22–24,34^. Although sustained deletion of ZNRF3/RNF43 eventually leads to HCC^22,29^, the underlying mechanism and the extent to which this mimics zonation disturbances seen in human CLDs remain unresolved.

Here, we address these gaps by integrating spatial, molecular, and lineage-tracing approaches across human CLD specimens, multiple murine disease models, and ZNRF3/RNF43-deficient livers. We show that expansion of β-catenin signaling from pericentral to periportal regions represents a conserved feature of diverse CLDs. We further dissect how this zonation collapse drives periportal hepatocyte proliferation, reprogramming, and tumor initiation, and demonstrate that the resulting ZNRF3/RNF43-deficient tumors represent a distinct metabolic and immunologic HCC subtype with clear parallels in human disease.

Together, our findings establish β-catenin–driven zonation disruption as a unifying, previously unrecognized mechanism that links chronic liver injury to malignant transformation, providing a conceptual framework for identifying high-risk patients and developing prevention strategies for CLD-associated HCC.

## Results

### Disruption of liver metabolic zonation is a common feature across diverse chronic liver diseases in human and mouse

Liver metabolic zonation ensures the spatial segregation of hepatocyte functions along the porto-central axis and is governed in part by WNT/β-catenin signaling. Although zonation disruption has been documented in metabolic dysfunction–associated steatohepatitis (MASH) in both mouse models and human liver^27,28^, whether zonation collapse represents a broader, etiology-independent feature of chronic liver disease (CLD) has remained unclear.

To address this question, we examined spatial distribution of CYP2E1, a canonical pericentral hepatocyte marker and direct β-catenin target, in human CLD specimens representing multiple etiologies, including MASH, chronic hepatitis C virus (HCV), hepatitis B virus (HBV), primary sclerosing cholangitis (PSC), primary biliary cholangitis (PBC), and alcoholic liver disease (ALD). In healthy liver, CYP2E1 expression was sharply restricted to the pericentral zone. In contrast, all CLD groups exhibited marked expansion of CYP2E1^+^ hepatocytes toward the periportal regions (**Figure 1A**), indicating loss of zonal confinement. Quantification revealed a significant increase in CYP2E1^+^ hepatic area from ∼55% in healthy livers to nearly 90-100% across all diseased livers (**Figure 1A-B**), establishing that pericentral gene expansion as a shared pathological feature of chronic liver injury.

**Figure 1.**
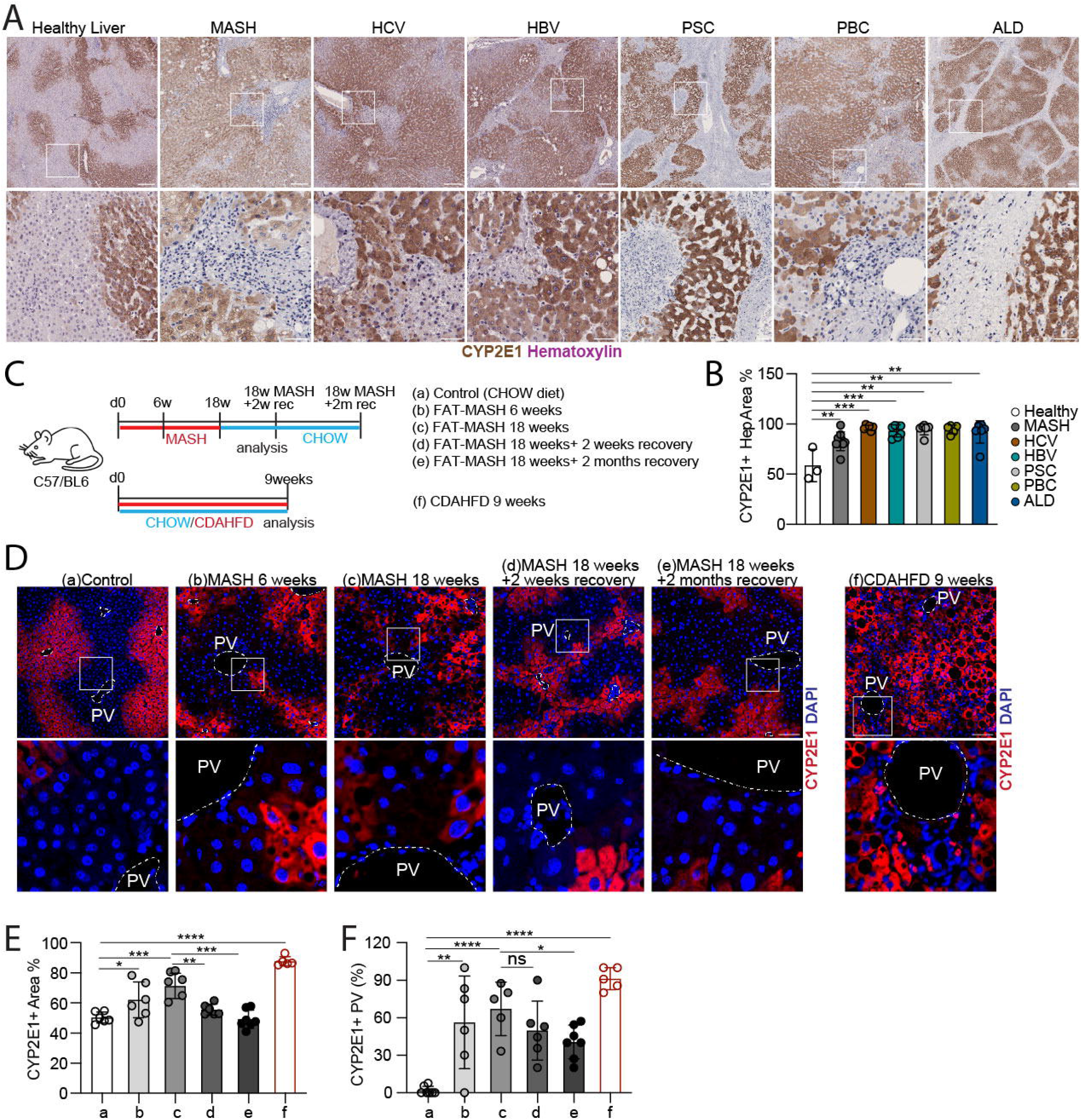
CYP2E1 zonation is altered across human liver diseases and in diet-induced mouse models. **(A)** IHC for CYP2E1 in human samples from healthy liver controls (n = 3), MASH (n = 7), HCV (n = 5), HBV (n = 6), PSC (n = 6), PBC (n = 5), and ALD (n = 6). Representative low-magnification images (4×, top row) and higher-magnification fields (20×, bottom row) are shown. **(B)** Quantification of CYP2E1-positive hepatocyte area (%) in the samples shown in (A), displayed in identical group order. **(C)** Schematics of mouse dietary experimental interventions. C57BL/6 mice were fed a FAT-MASH diet from day 0 to 18 weeks, followed by reversion to chow for 2 weeks or 2 months. Analysis timepoints include: (a) Chow control (n = 6), (b) FAT-MASH 6 weeks (n = 6), (c) FAT-MASH 18 weeks (n = 5), (d) FAT-MASH 18 weeks + 2-week recovery (n = 6), and (e) FAT-MASH 6 weeks + 2-month recovery (n = 7), (f) CDAHFD for 9 weeks. **(D)** IF staining for CYP2E1 and DAPI in liver sections from the timepoints shown in (C). Representative low-magnification images and higher-magnification views are shown. Portal and central veins are indicated as PV and CV, respectively. **(E)** Quantification of CYP2E1-positive area (%) for the samples shown in (D). **(F)** Quantification of CYP2E1-positive portal veins (%), defined as portal veins with ≥3 CYP2E1⁺ hepatocytes within the first three layers of the periportal region, for the samples shown in (D). Data are presented as mean ± s.d. with individual values overlaid. Statistical significance was determined by a two-tailed unpaired Student’s t-test for two-group comparisons or by a one-way ANOVA with *post hoc* correction for multiple-group comparisons. p < 0.05 (*), p < 0.01 (**), p < 0.001 (***), p < 0.0001 (****); ns, not significant. Scale bars: 500 µm (4×) and 100 µm (20×) (A, D).

To investigate whether zonation disruption is dynamic and reversible, we employed the well-established FAT (fibrosis and tumor)-MASH model, combining a high-fat Western diet with low-dose carbon tetrachloride, which robustly recapitulates fibrosis, advanced MASH, and early tumorigenesis^35,36^ (**Figure 1C**). Immunofluorescence analysis revealed progressive expansion of CYP2E1 from pericentral to periportal region during 6 -18 weeks of FAT-MASH feeding (**Figure 1D**, group a-c), accompanied by a significant increase in CYP2E1^+^ area (**Figure 1E**).

To test reversibility, we withdrew mice from FAT-MASH diet after 18 weeks and transitioned them to normal chow for either 2 weeks or 2 months. During this recovery period, total CYP2E1^+^ area gradually decreased toward baseline (**Figure 1D-E**, groups d-e); however, periportal aberrancy persisted. To assess zonation fidelity around the portal triad, we defined CYP2E1+ portal regions as those with ≥3 CYP2E1+ hepatocytes in the first three layers surrounding the portal vein (PV). Using this criterion, we found that 70% of portal regions became CYP2E1+ at 18 weeks of MASH, and notably, ∼50% remained CYP2E1+ even after 2 months of recovery (**Figure 1F)**, indicating that zonation defects only partially resolve and may be delayed relative to histological regression. We further validated these findings in an independent mouse MASH model using the choline-deficient amino acid-defined high-fat diet (CDAHFD), which similarly exhibited robust expansion of CYP2E1 toward periportal areas (**Figure 1D–F**, group f).

Since ZNRF3 and RNF43 act as negative regulators that restrict β-catenin activity signaling to the pericentral niche^23^, we hypothesized that hepatocyte-specific deletion of these E3 ligases would be sufficient to recapitulate CLD-associated β-catenin expansion in the absence of inflammation or fibrosis.

### Chronic liver disease-associated zone disruption is characterized by expansion of β-catenin activity from the pericentral to periportal zone and is recapitulated by ZNRF3/RNF43 deletion

To systematically and unbiasedly evaluate how chronic liver disease perturbs hepatic zonation, we quantified zonated gene expression along the periportal-pericentral (PP-PV) axis using established classification criteria^28^. Zonated genes were categorized into three major groups based on how their zonal enrichment changed relative to healthy liver: (1) less zoned, meaning genes that normally show strong enrichment in either the periportal or pericentral region begin to lose this spatial restriction and become more uniformly expressed across the lobule; (2) more zoned, in which already zone-enriched genes become even more sharply restricted to their original location, exaggerating the normal zonation pattern; and (3) swapped, where genes that are typically enriched in one zone (for example, pericentral) switch their enrichment pattern and become more highly expressed in the opposite zone (such as periportal), effectively inverting their spatial identity. Because hepatocyte zonation reflects opposing PP-PV and PV-PP, each category includes directional subtypes (**Figure 2A)**.

**Figure 2.**
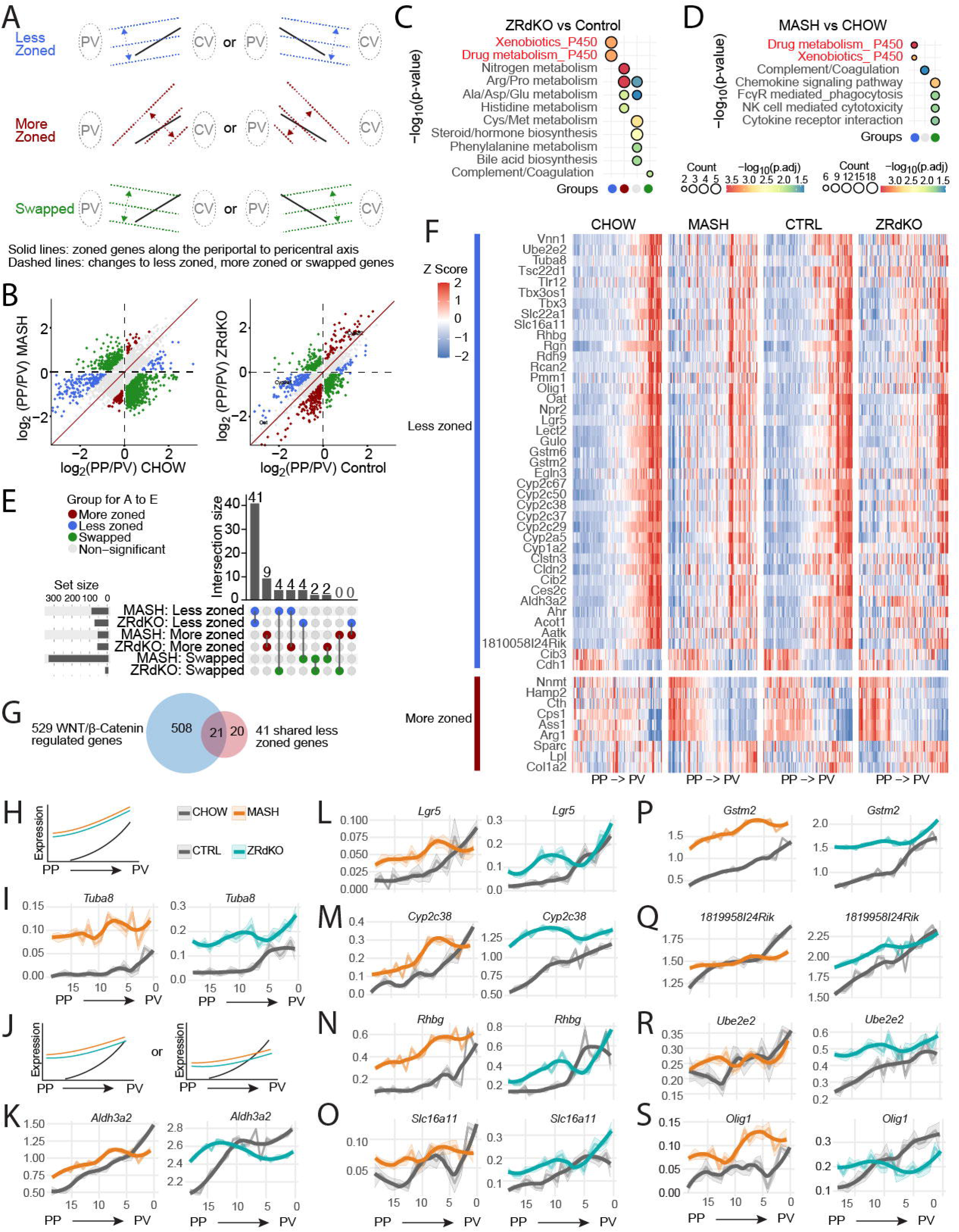
MASH and ZNRF3/RNF43-deficient livers share convergent WNT-driven zonation and transcriptional programs. **(A)** Schematic illustrating three classes of zonation changes: “Less zoned” (top row), “More zoned” (middle row), “Swapped” (bottom row). PV, portal vein; CV, central vein. **(B)** Scatter plots of gene expression between conditions. Left: log_2_ fold change of periportal/perivenous (PP/PV) expression in MASH versus CHOW. Right: log_2_(PP/PV) expression in ZRdKO versus Control. Vertical and horizontal dashed lines mark zero change. Genes are color-coded according to zonation category. **(C)** GO enrichment for zonation-shifted genes in ZRdKO versus Control. Ontology terms are shown on the y-axis. Dot color indicates log_10_(p-value), and dot size indicates gene count. Zonation changes categories are arranged along the x-axis. **(D)** GO enrichment for zonation-shifted genes in MASH versus CHOW. Ontology terms and visual encodings follow panel (C). **(E)** Intersection analysis of zonation-shift categories between MASH and ZRdKO. Rows represent the three zonation changes groups less zonated, more zonated, swapped in MASH and ZRdKO. Dots indicate presence in each set; connecting vertical lines denote intersections. The upper bar plot shows intersection sizes; the left bar plot indicates set sizes. **(F)** Heatmaps of zonation profiles in CHOW, MASH, CTRL, and ZRdKO samples. Transcriptional changes displayed as z-scores per gene are ordered along the periportal (PP)-to-perivenous (PV) axis. **(G)** Venn diagrams comparing WNT/β-catenin–regulated genes and the shared set of 41 less-zonated genes. The counts of exclusive and shared genes are indicated. (**H–J**) Schematic representation of spatial expression trends along the periportal–pericentral (PP→PV) axis. Grey lines denote baseline zonation profiles in CHOW and CTRL, while orange and turquoise lines represent corresponding trends in MASH and ZRdKO, respectively. The schematics illustrate genes that maintain conserved zonation patterns across all conditions as well as those displaying convergent or selectively altered periportal or pericentral expression in MASH and ZRdKO. (**I, K–S**) Spatial expression profiles of representative zonated genes corresponding to the schematic categories in (**H–J**). Expression is plotted along the PP to PV axis in arbitrary units, shown for CHOW vs MASH (left) and CTRL vs ZRdKO (right). Genes include *Tuba8* (**I**), *Aldh3a2, Lgr5, Cyp2c38, Rhbg, Slc16a11, Gstm2, 1819958I24Rik, Ube2e2*, and *Olig1* (**K–S**).

As both MASH and ZNRF3/RNF43 deletion exhibit expansion of WNT/β-catenin activity, we re-analyzed published spatial transcriptomics datasets from mouse MASH and from hepatocyte-specific ZNRF3/RNF43 deletion under homeostasis (**Suppl. Figure 1A-B**). Unbiased clustering readily distinguished hepatocyte groups corresponding to distinct zonal identities (**Suppl. Figure 1C**), and pseudotime ordering revealed a smooth periportal to midzonal and followed by pericentral continuum (**Suppl. Figure 1D-G**). Canonical zonation markers confirmed these assignments: periportal (*Ass1* and *Cyp2f2*) and pericentral (*Glul* and *Cyp2e1*). Both the MASH and ZNRF3/RNF43dKO datasets displayed robust expansion of pericentral identity (*Glul* and *Cyp2e1*) toward pericentral region toward periportal territories (**Suppl. Figure 1H-K**), matching histological observations (**Figure 1A**)^28^.

To visualize zonation changes more directly, we plotted the log₂(PP/PV) expression ratios for all zoned genes. In the MASH dataset, these ratios for chow versus MASH were plotted on the x- and y-axes, respectively, allowing us to classify genes into less zoned, more zoned, or swapped categories (**Figure 2B**, left). A parallel analysis comparing ZNRF3/RNF43dKO to control livers revealed similarly organized distributions (**Figure 2B**, right). Hallmark pathway enrichment analysis showed distinct biological signatures across more zoned, swapped, and unchanged genes. Notably, less zoned genes in both models were strongly enriched for detoxification and cytochrome P450 pathways (**Figure 2C-D**), implicating shared functional impairment of zonation-dependent hepatic metabolism.

To further dissect this shared signature, we identified 41 less zoned genes present in both datasets—representing over 50% of the less zoned cohort in ZNRF3/RNF43dKO (**Figure 2G**). Correlation metrics showed that these 41 genes clearly separated periportal and pericentral hepatocytes in healthy liver, but this zonal stratification collapsed in MASH and ZNRF3/RNF43dKO livers, where hepatocytes exhibited broad co-expression across normally restricted zonation markers (**Suppl. Figure 1L**), indicating global zonation collapse.

Among these shared 41 genes, 39 shifted unidirectionally from pericentral toward periportal expression, revealing the same directional expansion across both models (**Figure 2F**). This pervasive pericentral to periportal expansion suggests a conserved mechanism linking chronic liver injury to WNT/β-catenin dysregulation. Overlaying these genes with a curated set of 529 reported WNT**/**β-catenin–regulated genes confirmed that 21 are established β-catenin targets (**Figure 2G**), demonstrating that chronic disease and ZNRF3/RNF43 deletion converge on a common WNT/β-catenin-driven transcriptional program.

To better understand how individual genes changed across zones, we examined raw expression levels along the PP to PV axis. Among the 39 less-zoned genes that shifted consistently in both models, several genes including *Aatk, Ces2c, Clstn3, Npr2, Oat*, showed selective reduced expression in the pericentral zone while maintaining stable or only mildly reduced periportal expression (**Suppl. Figure 2A**). In contrast, *Tuba8* was the only gene uniformly upregulated across all liver zones in both MASH and ZNRF3/RNF43dKO (**Figure 2H-I**). A larger group of genes including *Tlr12, Pmm1, Gulo, Lect2, Rcan2, Vnn1, Tsc22d1, Rdh9* (**Suppl. Figure 2B**), along with tumor-associated genes such as *Aldh3a2, Gstm2, Lgr5, Rhbg, Slc16a11, Ube2e2, 1810058124Rlk, Cyp2c38, Olig1* (**Figure 2J-S**) retained their normal pericentral expression but exhibited substantial expansion into periportal zone^30,37–47^. Many of these are known β-catenin–responsive targets (Gstm2, Lect2, Lgr5, Rhbg, Tuba8, Ube2e2, Tsc22d1, Olig1), and several have documented roles in tumor suppression or tumor promotion, underscoring the potential oncogenic implications of their ectopic periportal activation.

Collectively, these findings demonstrate that periportal hepatocytes in chronic disease, and upon ZNRF3/RNF43 deletion acquire an aberrant hybrid identity, characterized by inappropriate activation of pericentral metabolic, detoxification, and tumor-associated gene programs. This ectopic activation reflects a profound, disease-associated loss of spatial hepatocyte specialization. This analysis reveals that β-catenin expansion from pericentral to periportal zones is the dominant shared feature between MASH and ZNRF3/RNF43 deletion

### Loss of ZNRF3/RNF43 induces β-catenin–dependent periportal hepatocyte proliferation and pericentral transcriptional reprogramming

ZNRF3 and RNF43 are E3 ubiquitin ligases that attenuate WNT/β-catenin signaling by promoting the degradation of Frizzled receptors^32,33^. Building on prior findings that loss of ZNRF3/RNF43 increases hepatocyte proliferation and converts periportal hepatocytes into pericentral-like states, we sought to more precisely define where these proliferating and reprogrammed hepatocytes arise within the lobule^22–24^.

To trace hepatocyte clone expansion across liver zones, we performed sparse lineage tracing using a low dose of AAV8-TBG-Cre to LSL-EGFP control mice or ZNRF3/RNF43; LSL-EGFP (ZNRF3/RNF43^ΔHepEGFP^) mice (**Figure 3A**). After two months, control livers contained only isolated GFP^+^ hepatocytes, consistent with minimal clonal expansion. In contrast, ZNRF3/RNF43-deleted livers exhibited large GFP⁺ hepatocyte clones, with a pronounced enrichment near periportal (PV) regions (**Figure 3B**). Quantification confirmed a marked increase in GFP⁺ hepatocytes in zone 1 (PV) and the periportal portion of zone 2 (PA), highlighting a strong periportal bias in proliferation (**Figure 3C**).

**Figure 3.**
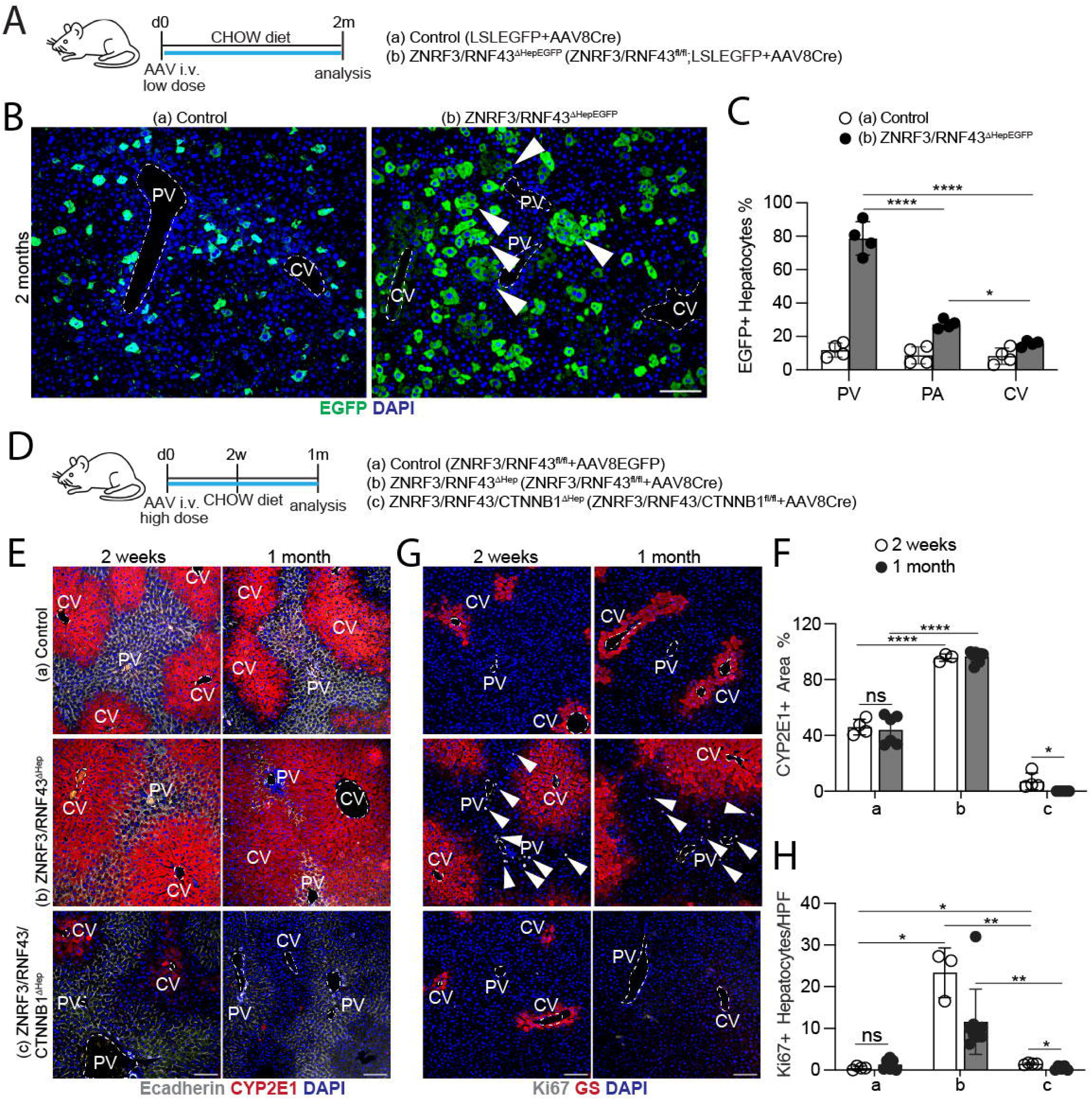
Hepatocyte-specific ZNRF3/RNF43 deletion drives periportal clonal expansion and induces WNT-mediated pericentral zone expansion. **(A)** Schematic of the low-dose AAV8Cre lineage-tracing strategy. Two experimental groups are included: (a) Control (LSLEGFP + AAV8Cre, n = 4) and (b) ZNRF3/RNF43^ΔHep-EGFP^ (ZNRF3/RNF43^fl/fl^; LSL-EGFP + AAV8Cre, n = 4). **(B)** Representative liver IF images for EGFP (green) and DAPI (blue) at 2 months post-injection for experimental groups in (A). **(C)** Quantification of EGFP^+^ hepatocytes in portal vein (PV), parenchyma (PA), and central vein (CV) regions of IF in (B). **(D)** Schematic of high-dose AAV8Cre KO strategy. Three experimental groups are included: (a) Control (ZNRF3/RNF43^fl/fl^ + AAV8-EGFP), (b) ZNRF3/RNF43^ΔHep^ (ZNRF3/RNF43^fl/fl^ + AAV8-Cre), and (c) ZNRF3/RNF43/CTNNB1^ΔHep^ (ZNRF3/RNF43/CTNNB1^fl/fl^ + AAV8Cre). n = 4 per group. **(E)** Representative liver IF images at 2 weeks and 1 month for each experimental group in (D), stained for E-cadherin, CYP2E1, and DAPI. **(F)** Quantification of CYP2E1^+^ area (% of total area) of IF in (E). **(G)** Representative liver IF images at 2 weeks and 1 month for each experimental group in (D), stained for Ki67, GS, and DAPI. **(H)** Quantification of Ki67^+^ hepatocytes per HPF from (G) at 2 weeks (white bars) and 1 month (dark grey bars) for each experimental group in (D). Arrows indicate Ki67^+^ cells. Data are presented as mean ± s.d. with individual values overlaid. Statistical significance was determined by a two-tailed unpaired Student’s t-test for two-group comparisons or by one-way ANOVA with appropriate *post hoc* correction for multiple-group comparisons. p < 0.05 (*), p < 0.01 (**), p < 0.0001 (****); ns, not significant. PV: portal vein, CV: central vein. Scale bars: 100 µm (B, E, G).

We hypothesized that β-catenin activity is required for both proliferation and zonal reprogramming. To test this, we induced hepatocyte-specific deletion of ZNRF3/RNF43 alone or in combination with β-catenin (CTNNB1) using AAV8-TBG-Cre (**Figure 3D**). As expected, ZNRF3/RNF43 deletion alone (group b) resulted in dramatic expansion of the pericentral marker CYP2E1 to nearly the entire lobule (**Figure 3E–F**). Similarly, GS (glutamine synthetase), normally restricted to the innermost 1–2 layers surrounding central veins, extending into midzonal and periportal territories (**Figure 3G**). This molecular reprogramming coincided with a substantial increase in Ki67⁺ proliferating hepatocytes, predominantly localized in the periportal zone at both 2 weeks and 1 month (**Figure 3G–H**). Strikingly, triple deletion of ZNRF3, RNF43, and CTNNB1 (group c) abolished these effects. CYP2E1 and GS expression were markedly reduced at 2 weeks and completely absent by 1 month, and the expansion of Ki67⁺ hepatocytes was entirely suppressed (**Figure 3E–H**, group c). These findings demonstrate that β-catenin is essential for both the proliferative surge and the periportal-to-pericentral transcriptional conversion induced by ZNRF3/RNF43 deletion.

Together, these findings establish that the aberrant expansion of β-catenin activity is the central driver of ZNRF3/RNF43 deletion-induced phenotypes. β-Catenin is required not only for periportal-biased proliferation but also for the acquisition of pericentral metabolic identity outside its normal niche. This provides mechanistic evidence that sustained β-catenin activation in the periportal zone disrupts zonal identity and primes hepatocytes for further pathological progression.

### ZNRF3/RNF43 deletion leads to spontaneous periportal-origin HCC

Recent studies suggest that hepatocyte metabolic zonation influences tumorigenic potential, raising the possibility that zone-specific reprogramming may shape where HCC originates. Although our prior work showing that long-term ZNRF3/RNF43 deletion cause spontaneous HCC^22^, the precise zonal origin was unclear. Because our earlier findings revealed robust periportal proliferation and pericentral-like reprogramming following ZNRF3/RNF43 loss, we hypothesized that ZNRF3/RNF43 deletion creates a pathological state in the periportal region from which HCCs arise.

To determine where tumors initiate, we performed 9-month clonal tracing using low-dose AAV8-TBG-Cre in LSL-EGFP controls and ZNRF3/RNF43^fl/fl^; LSL-EGFP mice (**Figure 4A**). In control livers, GFP⁺ hepatocytes appeared predominantly as single cells, doublets, or small clusters evenly distributed across lobule. In contrast, ZNRF3/RNF43-deleted livers exhibited large, expanding GFP⁺ hepatocyte clones predominantly localized near the periportal (PV) regions (**Figure 4B**). Quantification confirmed a dramatic enrichment of clones larger than 10 cells surrounding portal triads, indicating that periportal hepatocytes undergo sustained, long-term expansion when ZNRF3/RNF43 is absent (**Figure 4C**).

**Figure 4.**
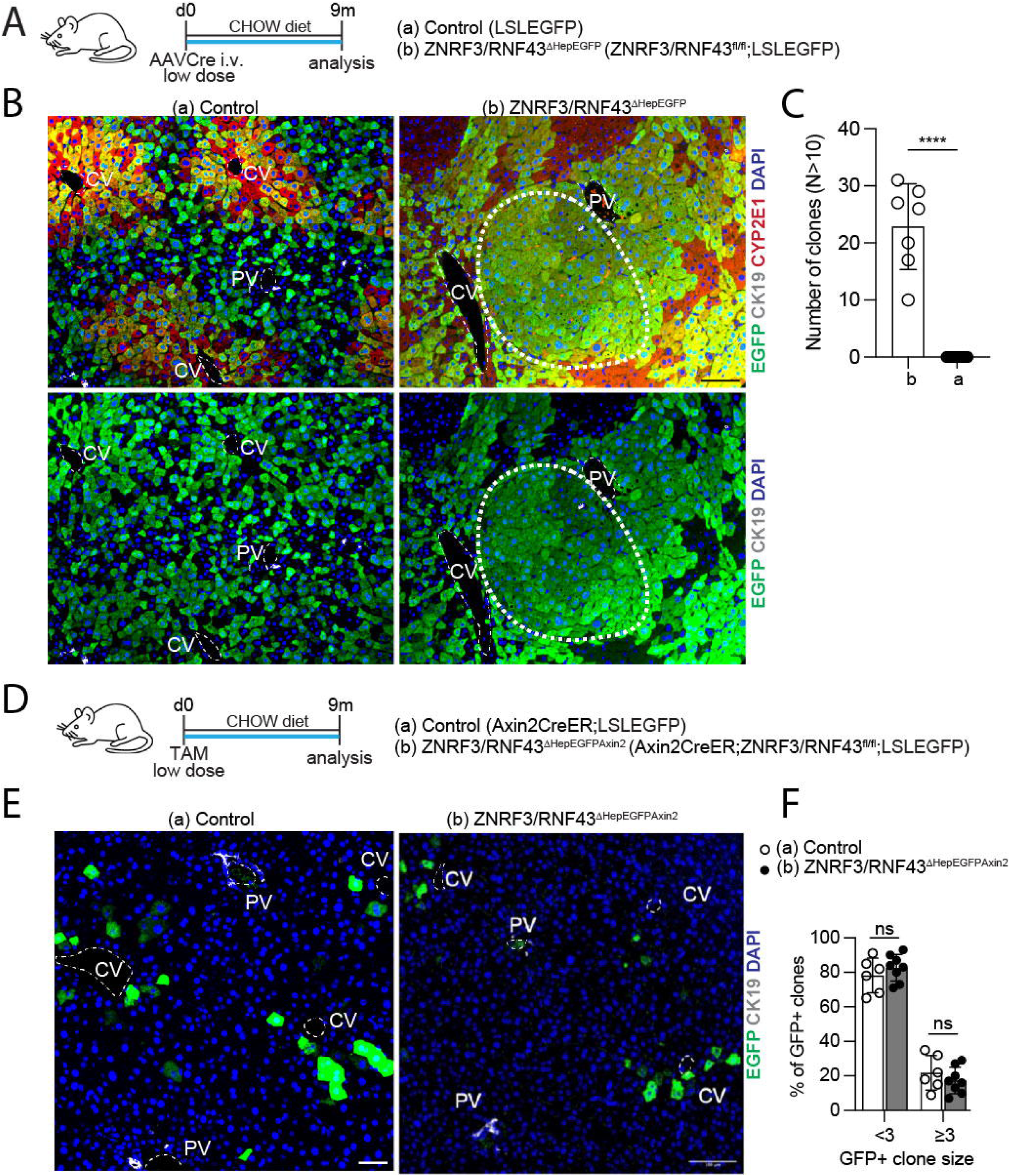
Periportal, but not pericentral, clonal expansion arises after hepatocyte-specific ZNRF3/RNF43 deletion in long-term lineage tracing. **(A)** Schematic of the low-dose AAVCre lineage-tracing experiment. Mice received a low i.v. dose of AAVCre at day 0 and were analyzed 9 months later. Experimental groups include: (a) Control (LSLEGFP) and (b) ZNRF3/RNF43^fl/fl^; LSLEGFP. n = 7 per group. **(B)** Representative liver IF images for EGFP, CK19, CYP2E1, DAPI, and EGFP, CK19, DAPI for groups in (A). **(C)** Quantification of GFP⁺ clones with size N > 10 for groups (a) and (b) from (B). **(D)** Schematic of the tamoxifen-inducible lineage-tracing experiment. Mice received a low dose of tamoxifen at day 0 and were analyzed 9 months later. Experimental groups: (a) Control (Axin2CreER; LSLEGFP – n = 6) and (b) ZNRF3/RNF43^ΔHepEGFPAxin^^2^ (ZNRF3/RNF43^fl/fl^; Axin2CreER; LSLEGFP - n = 8). **(E)** Representative IF images for the groups in (D), stained for EGFP (green), CK19 (grey), and DAPI (blue). **(F)** Quantification of % of GFP⁺ clone distribution by size: < 3 cells and ≥ 3 cells for experimental groups in (D). Data are presented as mean ± s.d. with individual values overlaid. Significance was determined using a two-tailed unpaired Student’s t-test for pairwise comparisons. Significance levels: p < 0.0001 (****); ns, not significant. PV: portal vein, CV: central vein. Scale bars: 100 µm (B, E)

Although early preneoplastic lesions were visible at 9 months, their size and mixed zonal composition made the exact cell-of-origin difficult to determine. Thus, an additional lineage-tracing approach was required to determine whether pericentral hepatocytes also contribute to HCC formation.

Using AXIN2CreER, which labels exclusively pericentral hepatocytes following low-dose tamoxifen, we tracked the fate of pericentral cells over 9 months in control (AXIN2CreER; LSL-EGFP) and ZNRF3/RNF43-deleted (ZNRF3/RNF43^fl/fl^; AXIN2CreER; LSL-EGFP) mice. In both groups, AXIN2-labeled hepatocytes remained confined to the pericentral region and persisted only as isolated cells or small clusters (**Figure 4D**). Crucially, no large AXIN2-derived clones were detected in ZNRF3/RNF43-deleted livers, demonstrating that pericentral hepatocytes do not expand clonally and do not contribute to tumor initiation in this model (**Figure 4F**).

Together, these complementary lineage-tracing approaches demonstrate that spontaneous HCCs arising after ZNRF3/RNF43 deletion originate from the periportal zone, where β-catenin activity becomes ectopically and persistently elevated. These results firmly establish that ZNRF3/RNF43-mediated zonation disruption creates a periportal tumor-permissive state, linking aberrant β-catenin expansion to the earliest steps of hepatocarcinogenesis.

### HCCs arising from ZNRF3/RNF43 deletion exhibit distinct zonal, metabolic and immune signatures

Having established that ZNRF3/RNF43 deletion induces spontaneous HCCs originating from the periportal zone (**Figure 4**), we next sought to define the molecular identity of these tumors and determine how they compare with classical oncogene-driven HCC models. Specifically, we investigated whether the zonal origin of ZNRF3/RNF43-deficient tumors imprints characteristic transcriptional, metabolic, and immunologic features.

We performed bulk RNA sequencing on ZNRF3/RNF43-deficient tumors (ZRdKO tumors), matched non-tumor liver tissues from the same mice (ZRdKO normal), and ZNRF3/RNF43 wild-type livers (ZRdKO wild-type). These datasets were analyzed alongside two published murine HCC cohorts, including DEN-induced tumors (DEN tumors)^22^ and genetically engineered tumors driven by CTNNB1 or MYC^48^.

Multidimensional scaling analysis revealed that all non-tumor liver samples, regardless of genotype, clustered tightly together. In contrast, ZRdKO tumors formed a distinct and cohesive cluster, positioned closest to DEN tumors but clearly separated from both CTNNB1-driven and MYC-driven tumors (**Figure 5A**). This demonstrates that ZRdKO tumors represent a unique molecular class not captured by canonical oncogenic drivers.

**Figure 5.**
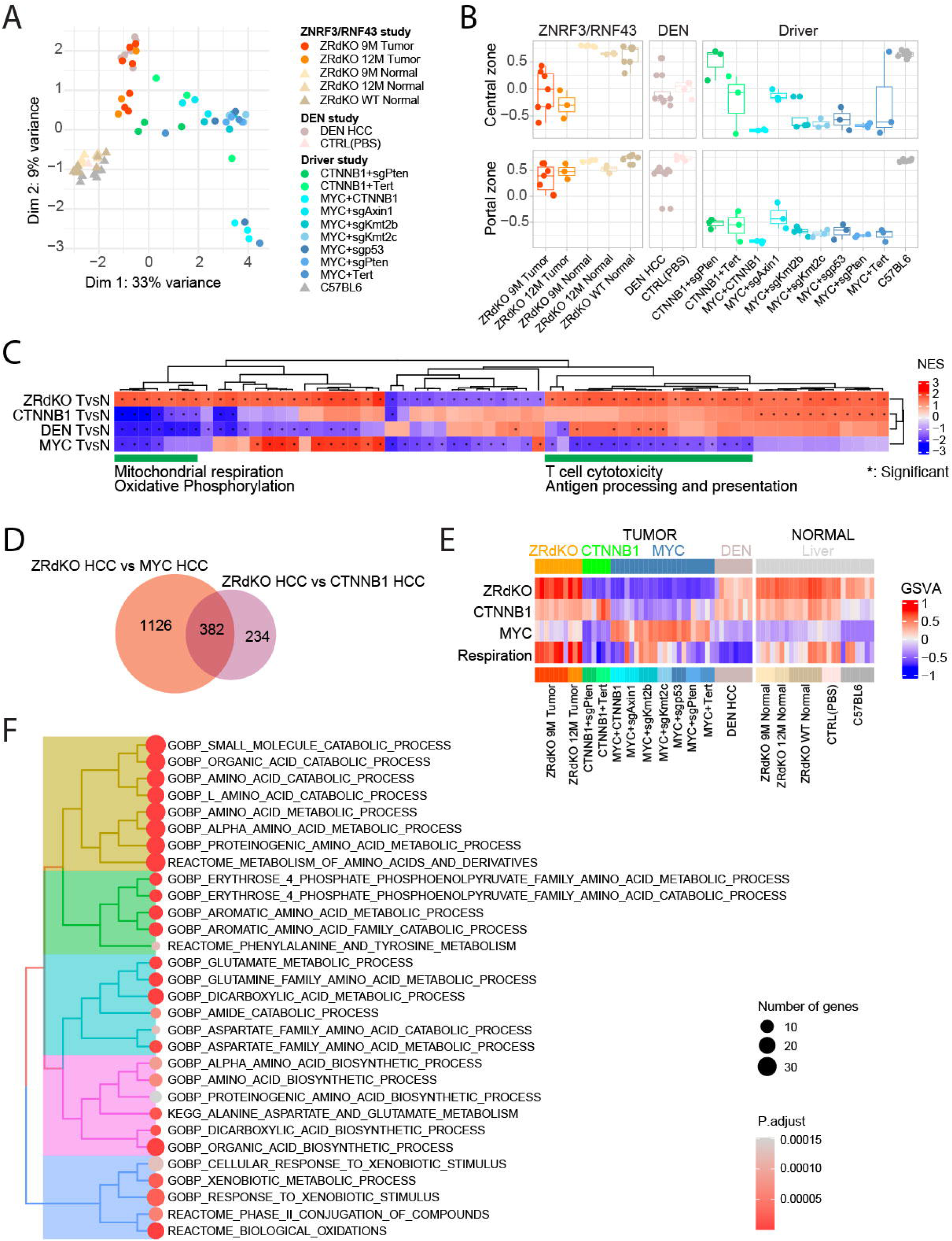
ZNRF3/RNF43-deficient HCCs form a portal-biased, respiration-high subset among mouse liver tumor models. **(A)** Multidimensional scaling plot (MDS) of bulk RNA-seq from ZRdKO tumors, matched adjacent non-tumor liver tissue, DEN-induced HCCs, and MYC- or CTNNB1-driven tumors. Distances on the plot approximate the typical log2 FC between the samples. **(B)** Box plots of GSVA-derived zonation scores for the same samples shown in (A). Portal (zone 1) and central (zone 3) signatures are indicated. **(C)** Heatmap of GSEA normalized enrichment scores for tumor-versus-normal comparisons across ZRdKO, CTNNB1, DEN, and MYC models. Pathways and model contrasts are clustered hierarchically. Asterisks denote statistically significant enrichments. **(D)** Venn diagrams showing overlap of differentially expressed genes (tumor versus normal) between ZRdKO tumors and MYC-driven tumors, and between ZRdKO tumors and CTNNB1-driven tumors. **(E)** Heatmap of GSVA scores for ZRdKO, CTNNB1, MYC, and respiration signatures across all tumor and non-tumor liver samples, with columns grouped by model and by tumor versus normal status. Respiration signature was computed using the leading-edge genes from the pathways highlighted in green in (C). **(F)** Dot-plot representation of functional enrichment analysis of the ZRdKO HCC gene signature. Dot size corresponds to the number of genes associated with each ontology term, and dot color reflects adjusted P values. Ontology terms are hierarchically clustered.

Because ZRdKO tumors arise from the periportal region, we assessed whether they retain transcriptional feature of their cell-of-origin. Gene set variation analysis (GSVA) using zonation signatures showed that healthy livers from all cohorts displayed strong periportal and pericentral transcriptional identities, whereas ZRdKO tumors demonstrated a marked loss of pericentral identity while maintaining high periportal signatures (**Figure 5B; Suppl. Figure 3A–B**). Interestingly, DEN tumors partially mirrored this pattern. In contrast, CTNNB1-driven tumors and MYC-driven tumors exhibited highly variable zonation profiles but consistently lost periportal signatures, making a high periportal signature a distinguishing feature of ZRdKO tumors.

To identify pathways altered during tumorigenesis, we compared ZRdKO tumors with matched non-tumor liver using gene-set enrichment analysis (GSEA). ZRdKO tumors demonstrated significant upregulation of oxidative phosphorylation, mitochondrial ATP synthesis and electron transport chain activity (**Figure 5C; Suppl. Figure 3C**). These pathways collectively indicate a state of elevated mitochondrial respiration, which stands in contrast to the glycolytic preference commonly observed in many cancers. ZRdKO tumors also showed enrichment of amino-acid catabolism and detoxification pathways, consistent with the metabolic programs normally enriched in periportal hepatocytes (**Figure 5F)**. Unlike MYC-driven or CTNNB1-driven HCCs, which typically exhibit immune-cold phenotypes, ZRdKO tumors displayed increased T cell cytotoxicity, enhanced leukocyte activation, and upregulation of antigen presentation pathways (**Figure 5C; Suppl. Figure 3C**). This suggests that ZRdKO tumors maintain or even amplify immune visibility, raising the possibility that periportal-origin HCCs could respond more favorably to immunotherapies.

Whole-exome sequencing of ZRdKO tumors revealed sparse and largely non-recurrent mutations, with minimal overlap with known HCC driver genes (**Suppl. Figure 3D-E**). Tumor mutational burden remained low, even in 12-month tumors (**Suppl. Figure 3D**). This indicates that metabolic and transcriptional reprogramming—not genetic mutation—drives tumor evolution in the ZNRF3/RNF43-deficient context.

By comparing ZRdKO tumors with CTNNB1- and MYC-driven tumors, we identified a set of 382 genes uniquely upregulated in ZRdKO tumors (**Figure 5D**). Functional enrichment showed that these genes are strongly associated with pathways involved in mitochondrial respiration, amino acid catabolism, detoxification, and broader metabolic processes (**Figure 5E**). These transcriptional programs echo the metabolic identity of periportal hepatocytes and reinforce the concept that the zonal origin of ZRdKO tumors leaves a lasting metabolic imprinting on tumor phenotype.

Together, these findings establish that ZNRF3/RNF43 deletion produces a molecularly distinct HCC subtype characterized by preserved periportal identity, enhanced mitochondrial metabolism, and enhanced immune activity, while lacking recurrent classical cancer-driver mutations. These unique features distinguish ZRdKO tumors from CTNNB1-driven, MYC-driven tumors and highlight periportal-origin, β-catenin-expanded tumors as a biologically coherent and potentially therapeutically responsive HCC subclass. Having defined the molecular features of ZRdKO tumors, we next asked whether a subset of human HCCs shares these characteristics.

### ZNRF3/RNF43dKO tumors represent a subpopulation of human HCCs

ZNRF3/RNF43 deletion releases β-catenin activity into the periportal zone and rewires periportal hepatocyte metabolism, ultimately enabling tumor initiation at this location. Because chronic liver disease also exhibits periportal expansion of β-catenin programs and tumor-associated genes, we hypothesized that a subset of human HCCs shares transcriptional features with ZRdKO tumors.

To identify ZRdKO-like tumors in human datasets, we applied the ZRdKO tumor signature to TCGA^49^ and MERiC^50^ HCC cohorts and defined ZRdKO-like HCCs as tumors with the top 25% of ZRdKO signature scores and simultaneously within the top 25% for differential enrichment relative to CTNNB1- and MYC-driven signatures. This approach identified a distinct group of ZRdKO-like HCCs, with all remaining tumors classified as other HCCs (**Figure 6A**).

**Figure 6.**
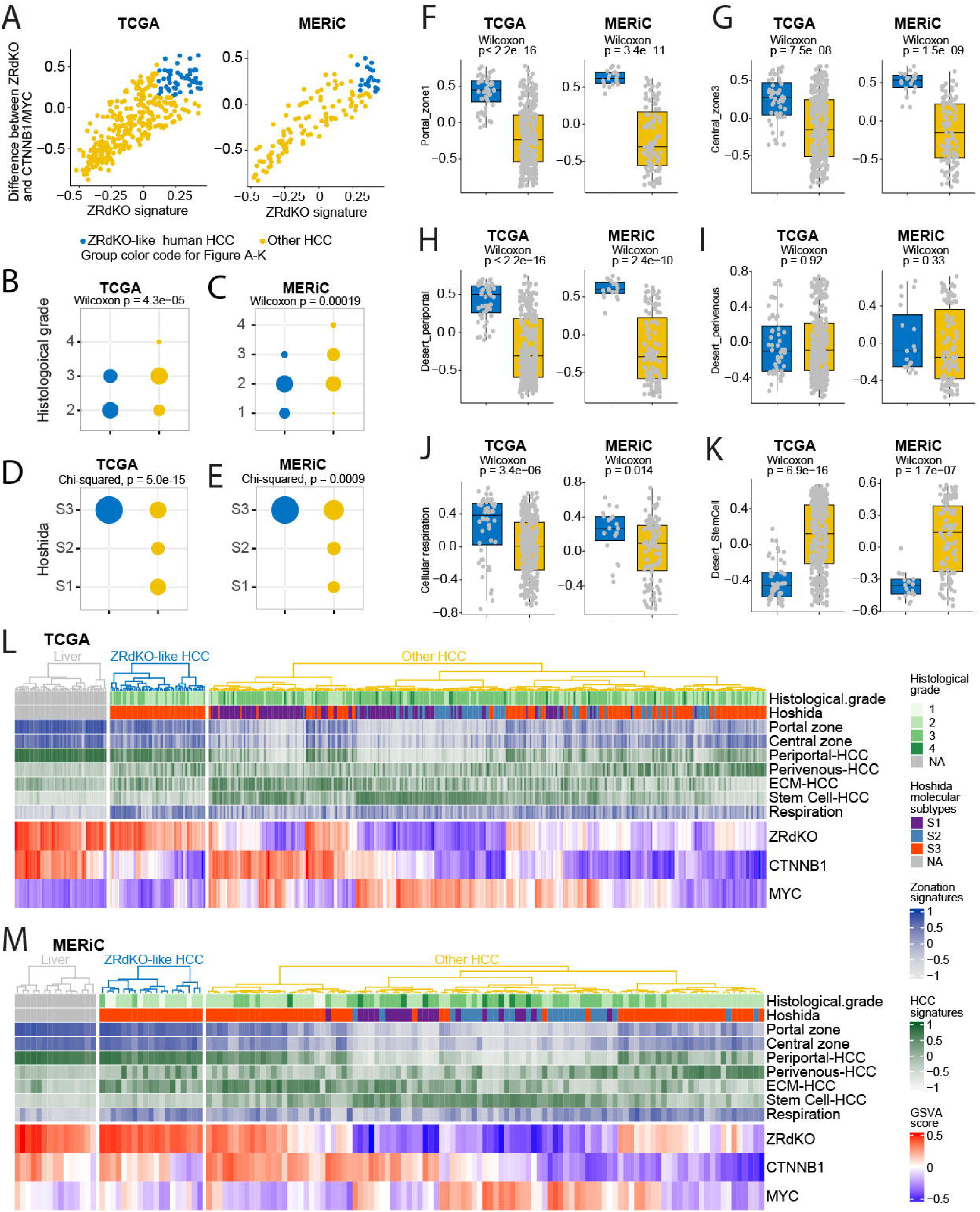
ZRdKO-like human HCCs define a portal-enriched, low-grade metabolic subset in TCGA and MERiC cohorts. **(A)** Scatter plots showing ZRdKO-like classification in TCGA (left) and MERiC (right) based on the difference between ZRdKO and CTNNB1/MYC signature scores. **(B-C)** Dot plots comparing histological grade between ZRdKO-like and Other HCC in TCGA (B) and (C). Dot size indicates the fraction of ZRdKO-like HCC or Other HCC. Wilcoxon rank-sum P values are shown. **(D-E)** Dot plots showing distribution of Hoshida molecular subclasses (S1-S3) in ZRdKO-like and Other HCC in TCGA (D) and MERiC (E). Dot size indicates the fraction of ZRdKO-like HCC or Other HCC. Chi-squared P values are shown. **(F-K)** Box plots comparing GSVA scores between ZRdKO-like and Other HCC in TCGA (left) and MERiC (right) for: Portal zonation (F), Central zonation (G), Periportal HCC (H), Perivenous HCC (I), Respiration (J), and Stem cell HCC (K) signatures. Wilcoxon rank-sum P values are shown. (L–M) Heatmaps of histological, molecular and transcriptional features for TCGA (L) and MERiC (M) cohorts, grouped into non-tumor liver, ZRdKO-like HCC, and Other HCC. For TCGA, the non-tumor group corresponds to adjacent liver tissue from HCC patients; for MERiC, it corresponds to healthy liver from non-HCC individuals. Rows include histological grade, Hoshida molecular subtype, zonation signatures, HCC signatures, the respiration signature, and GSVA scores for ZRdKO, CTNNB1 and MYC. Columns within each subgroup are hierarchically clustered.

ZRdKO-like HCCs showed reduced ZNRF3 and RNF43 expression compared to other HCCs, although statistical significance in MERiC cohort was limited by sample size (**Suppl. Fig. 4D-E**). Relative to other HCCs, ZRdKO-like HCCs exhibited lower histological grade (**Figure 6B-C**) and were strongly enriched for the Hoshida S3 subtype, consistent with the low-grade, metabolic specialized phenotype of ZRdKO mouse tumor (**Figure 6D-E**, **Figure 5F**). Importantly, they did not differ in TP53, ARID1A, or CTNNB1 mutation status (**Suppl. Fig. 4A-C**), indicating that their identity reflects a transcriptional state rather than genetic alterations.

Consistent with the mouse findings, ZRdKO-like HCCs displayed higher periportal and pericentral zonation signatures (**Figure 6F-G**) and a higher periportal-HCC signature but not pericentral-HCC signature (**Figure 6H-I**). These patterns support a strong transcriptional resemblance to periportal-origin ZRdKO tumors. ZRdKO-like HCCs also showed significant enrichment of mitochondrial respiration pathways **(Figure 6J)**, matching the elevated oxidative metabolism observed in mouse ZRdKO tumors. Furthermore, ZRdKO-like HCCs exhibited significantly lower Desert stem-cell scores (**Figure 6K**), with no differences in Desert ECM signatures (**Suppl. Fig. 4F**), reinforcing their low-grade phenotype.

Across both human cohorts, ZRdKO signature scores correlated strongly with portal-zone and periportal-HCC signatures (r > 0.8; **Suppl. Fig. 4I-J**). Despite these strong correlations, gene overlap among ZRdKO, Desert periportal-HCC, and portal-zone signatures was limited (**Suppl. Fig. 4H**), demonstrating that these signatures represent convergent but mechanistically distinct aspects of periportal biology.

Hierarchical clustering based on ZRdKO-, CTNNB1-, and MYC-derived gene signatures further confirmed that ZRdKO-like HCCs form a coherent molecular subgroup in both TCGA and MERiC (**Figure 6L-M**). This subgroup is characterized by low grade, S3 molecular status, preserved periportal/pericentral zonation, enrichment for the periportal-HCC subclass, and elevated mitochondrial respiration.

Collectively, this analysis reveals that approximately 5-10% of human HCCs share the zonal, metabolic, and transcriptional properties of ZRdKO tumors. This work identifies a previously unrecognized subgroup of human HCC defined by periportal β-catenin mislocalization, a zonation-disrupting event that reprograms hepatocyte identity and metabolic state to generate a distinct, low-grade periportal tumor phenotype.

## Discussion

Metabolic zonation along the porto–central axis enables hepatocytes to perform distinct functions, and a graded WNT/β-catenin signaling landscape is central to maintaining this organization^22–24^. During acute injury, β-catenin activation is briefly induced across zones but rapidly resolves once negative feedback becomes re-engaged^21^. This transient, spatially confined activation supports regeneration while preserving zonal architecture^21^. In chronic liver disease, this regulatory system fails^29^. Using spatial transcriptomics and immunohistochemistry across multiple human CLD etiologies including MASLD/MASH, viral hepatitis, cholangiopathies, and alcohol-associated liver disease, we demonstrate that sustained, spatially aberrant β-catenin expansion from pericentral to periportal territories is a conserved pathological feature independent of disease etiology. This zonation collapse is not merely correlative; hepatocyte-specific deletion of ZNRF3/RNF43, which enforces sustained periportal β-catenin activity in the absence of inflammation or fibrosis, directly recapitulates this expansion and drives spontaneous periportal-origin HCC after long-term exposure.

Our findings establish a mechanistic framework in which chronic injury leads to sustained β-catenin zonation disruption induced periportal reprogramming, which contributes to malignant transformation, and represents a previously unrecognized pathway to hepatocarcinogenesis. Critically, this pathway operates through transcriptional and metabolic reprogramming rather than classical mutational drivers. ZNRF3/RNF43-deficient tumors harbor minimal recurrent mutations yet exhibit profound metabolic rewiring, preserved periportal transcriptional identity, enhanced mitochondrial respiration, and immune competence, features that distinguish them from CTNNB1- or MYC-driven HCCs and that correspond to approximately 5-10% of human HCCs. Together, these results reveal β-catenin-driven zonation disruption as a unifying, etiology-independent mechanism linking chronic liver injury to malignant transformation.

Our spatial transcriptomic analyses reveals that chronic injury and ZNRF3/RNF43 deletion converge on a shared transcriptional mechanism. Among 41 "less-zoned" genes that lose spatial restriction in both conditions, 21 are established β-catenin targets, and many genes including *Gstm2* and others have documented roles in tumorigenesis. Crucially, these genes are not globally upregulated; rather, they become inappropriately expressed in the periportal zone where they are normally silenced. This spatial dysregulation would be invisible to bulk transcriptomic analyses, underscoring that zonal context, not absolute expression level, determines oncogenic potential.

This concept is reinforced by recent studies demonstrating that hepatocyte zonal identity fundamentally shapes tumorigenic susceptibility. Work from the Zhu laboratory showed that in a *Ctnnb1/Arid2*-driven model, HCCs arise preferentially from zone 3, where pericentral genes like *Gstm2* and *Gstm3* are required for tumor initiation^30^. The Sansom laboratory demonstrated that GLUL⁺/LGR5⁺ pericentral hepatocytes resist CTNNB1/MYC-induced transformation^31^. Our findings extend this framework by showing that β-catenin expansion into the periportal region rather than simply increased β-catenin amplitude, is the critical event conferring tumor susceptibility. Upon ZNRF3/RNF43 deletion, β-catenin is activated uniformly across all zones, yet only periportal hepatocytes undergo sustained clonal expansion and give rise to HCC.

Importantly, although many periportal clones expand following ZNRF3/RNF43 deletion, only a minority progress to tumors, indicating that additional selective pressures beyond initial β-catenin-driven proliferation are required for malignant transformation. The identity of these secondary factors—whether metabolic, epigenetic, or microenvironmental—remains to be defined.

ZNRF3/RNF43-deficient tumors represent a molecularly distinct HCC subtype defined by three key features: preserved periportal identity, enhanced mitochondrial metabolism, and immune competence. Unlike classical HCC models driven by oncogenic mutations, ZRdKO tumors lack recurrent driver alterations even after one year of evolution, suggesting that metabolic and transcriptional state, rather than genetic lesions, sustains tumor growth. Their elevated oxidative phosphorylation and amino acid catabolism stands in stark contrast to the glycolytic preference typical of many cancers and likely reflects their periportal origin^51^, where these metabolic pathways are normally enriched.

The immune landscape of ZRdKO tumors is particularly notable. Whereas CTNNB1- and MYC-driven HCCs typically exhibit immune-cold phenotypes with minimal T-cell infiltration^52–54^, ZRdKO tumors maintain signatures of T-cell cytotoxicity, leukocyte activation, and antigen presentation. This immune visibility suggests that periportal-origin, β-catenin-expanded HCCs may respond more favorably to immune checkpoint blockade than other HCC subtypes. The identification of a corresponding ZRdKO-like subset comprising 5-10% of human HCCs, characterized by low histological grade, Hoshida S3 molecular classification, preserved zonation signatures, and elevated mitochondrial respiration, provides a framework for prospectively identifying patients who might benefit from immunotherapy or metabolism-targeted interventions.

The unique metabolic dependencies of ZRdKO tumors also present therapeutic opportunities. Their reliance on oxidative phosphorylation, rather than glycolysis, suggests vulnerability to mitochondrial inhibitors, while their enrichment of amino acid catabolic pathways points to potential metabolic liabilities distinct from other HCC subclasses. Furthermore, the absence of classical driver mutations implies that these tumors may be less amenable to targeted therapies aimed at specific oncogenic lesions but more responsive to strategies that disrupt their metabolic or zonal identity.

From a clinical perspective, these findings have immediate implications for HCC risk stratification in CLD patients. The observation that zonation disruption is conserved across etiologically diverse CLDs suggests that spatial assessment of β-catenin target genes, such as CYP2E1 expansion into periportal regions, could serve as a biomarker for elevated HCC risk independent of traditional clinical parameters like fibrosis stage. Patients exhibiting persistent periportal CYP2E1 expression despite histological improvement may represent a high-risk population warranting intensified surveillance or preventive intervention.

Although our findings establish β-catenin expansion as essential for early periportal proliferation and reprogramming, evaluating its requirement during long-term tumorigenesis remains technically challenging. Global β-catenin deletion causes profound metabolic dysfunction, including lipid accumulation, bile acid dysregulation, and impaired cell adhesion, confounding interpretation of its tumor-suppressive role in this context^55,56^. Future studies employing periportal zone-specific β-catenin inhibition or selective blockade of its nuclear transcriptional function will be necessary to definitively assess whether sustained β-catenin activity is required throughout the full trajectory from zonation disruption to malignant transformation^52^.

Although β-Catenin expansion from pericentral to periportal zone is a key feature of MASH and ZNRF3/RNF43 deletion^22,28^, and multiple identified less zoned genes expanded to periportal zone are β-Catenin targets and has been reported to be involved in tumorigenesis, the direct regulation of these genes involved in β-Catenin is to be defined by exploring the incorporation of β-Catenin with potential distinct transcriptional factors for DNA binding. Interestingly, ZNRF3/RNF43 deletion and MASH livers showed an expansion of β-catenin rather than an upregulation of its activity across all liver zones. This is reflected by the selective expression of β-catenin targets in specific zones, and such selective zonal regulation can derive from both extrinsic signaling and intrinsic chromatin structure.

Finally, β-Catenin activation itself has been shown to be not sufficient to induce HCCs, and secondary drivers or factors are required for the potential malignant transformation^57^. ZNRF3/RNF43 deletion elevated multiple metabolic pathways including fatty acid, lipid and bile acid metabolism while tumor associated pathways are not detected^29^. Although ZNRF3/RFN43 deletion drives β-Catenin activation at the periportal zone and the formation of periportal HCCs, how other factors and pathways contribute to tumorigenesis at certain timeline is not clear. A systematic longitudinal study to map the metabolic reprograming and tumorigenesis process at the spatial level is important to understand the key steps of pathogenesis and to provide insights into the safety window for applying regenerative medicine.

Our findings reveal β-Catenin activity expansion from their original pericentral zone to periportal region is etiology-independent mechanism shared in various chronic liver diseases and provide a tractable genetic model ZNRF3/RNF43 deletion to mechanistically dissect this pathway. Tracing hepatocytes across liver zones upon ZNRF3/RNF43 deletion revealed the increased periportal hepatocyte proliferation followed by the origin of HCC from the periportal zone, suggesting a shared tumorigenesis mechanism may exist in chronic liver diseases. The identification of a corresponding ZRdKO-like subset in human HCC, defined by preserved zonation, elevated mitochondrial metabolism, immune competence, and low-grade histology, establishes molecular criteria for patient stratification and suggests targeted therapeutic approaches.

From a preventive oncology perspective, monitoring zonation fidelity in CLD patients could enable early identification of individuals at elevated HCC risk. Simple immunohistochemical assessment of CYP2E1 or GS expansion into periportal regions on surveillance liver biopsies represents a readily implementable biomarker strategy. Patients exhibiting persistent zonation disruption despite treatment of their underlying liver disease may benefit from intensified imaging surveillance or consideration of preventive interventions targeting β-catenin signaling or its downstream metabolic programs.

For patients with established ZRdKO-like HCCs, the unique metabolic and immune features of this subtype suggest distinct therapeutic vulnerabilities. The preserved immune competence, in contrast to immune-cold CTNNB1/MYC-driven tumors, positions this subset as a high-priority candidate for immune checkpoint blockade. The reliance on oxidative phosphorylation and amino acid catabolism points to targetable metabolic dependencies that could be exploited with mitochondrial inhibitors or pathway-specific metabolic blockers.

In summary, this study establishes zonation disruption—specifically, sustained periportal β-catenin expansion—as a unifying, mechanistically grounded pathway from chronic liver injury to HCC, opening new avenues for risk stratification, early detection, and precision therapy in chronic liver disease-associated hepatocellular carcinoma.

## Acknowledgements

We thank Jade Tam, Sophia Wang for their technical support. This study was supported by Icahn School of Medicine at Mount Sinai.

## Author contributions

Conceptualization, T.S., F.D.T., S.B.Y. L.Y., C.K.Y.N; Methodology, F.D.T., S.B.Y. S.R., B.C., D.S., J.C., C.K.Y.N., J.I.A., J.S.T., S.M., A.B., L.L., L.Y., T.S.; Formal analysis, F.D.T., S.B.Y. S.R., B.C., D.S., J.C., C.K.Y.N., J.I.A., S.M., L.Y, T.S.; Investigation, F.D.T., S.B.Y. S.R., J.S.T., J.I.A., S.M., L.Y, T.S.; Supervision, L.Y., C.K.Y.N., J.S.T., S.M., T.S.; Writing – original draft, S.B.Y., F.D.T., L.Y., C.K.Y.N., S.M., T.S..

## Materials availability

This study did not produce any new or unique reagents.

## Data and software availability

Bulk RNA-seq data are available at the Gene Expression Omnibus under the accession (pending). Whole exome sequencing data are available at the NIH Sequence Read Archive under the designated BioProject ID (pending). All software and algorithms use for data analyses are previously published and appropriately referenced. Further correspondence and requests for resources and reagents should be addressed to Tianliang Sun.

## METHODS

### Ethics statement

All animal procedures were performed in accordance with institutional guidelines and approved by the Institutional Animal Care and Use Committee (IACUC) of the Icahn School of Medicine at Mount Sinai (ISMMS). Mice were monitored regularly for health and well-being and were provided ad libitum access to food and water. Humane endpoints were defined in accordance with ISMMS animal care policies.

### Human samples

Formalin-fixed, paraffin-embedded (FFPE) human liver tissue was obtained from Biorepository Core at Icahn School of Medicine at Mount Sinai including healthy controls and patients with metabolic dysfunction–associated steatohepatitis (MASH), chronic hepatitis C virus infection (HCV), chronic hepatitis B virus infection (HBV), primary sclerosing cholangitis (PSC), primary biliary cholangitis (PBC), and alcohol-associated liver disease (ALD). Tissue handling, fixation, embedding, and sectioning were performed according to standard pathology protocols, as detailed below.

### Mouse Husbandry and specimens’ collection

Mice were housed in the Icahn Building Mouse Facility at ISMMS under a 12-hour light/dark cycle in filtered cages with nesting enrichment. Breeding cages were limited to two or three mice, and experimental cages to a maximum of five mice. Single housing was used only when necessary (i.e. aggressive behavior). Mice were permanently identified at weaning (postnatal day 21) using the GEMS (Genetically Engineered Mouse Subcommittee) ear-notching system, encoding numerical IDs via predefined left/right ear notch patterns. In some experiments, alternative identification methods (ear punches or numbered clips) were used, employing sterile tools and performed by trained personnel. Age-matched adult male and female mice were randomly assigned to experimental groups in balanced numbers, unless otherwise stated.

Animals were monitored daily for health, body weight, and signs of distress. At experimental endpoints, or when humane euthanasia criteria were met, mice were euthanized by CO₂ asphyxiation followed by cervical dislocation, in accordance with IACUC guidelines, blood was collected from the vena cava for serum isolation, and livers were dissected. The caudate lobe was snap-frozen in liquid nitrogen for molecular analyses; standardized regions of left, median, and right lobes were fixed in 10% neutral-buffered formalin (ThermoFisher, Cat# 23-245685) and embedded in OCT (ThermoFisher, Cat# 23-730-571) and processed as described below.

### Tissue processing

Liver tissue was fixed in 10% neutral-buffered formalin (ThermoFisher, Cat# 23-245685) for 48 h, trimmed after 24 h, and transferred to 50% ethanol prior to paraffin embedding using a VIP tissue processor. Sections (3 µm) were cut onto glass slides. For frozen sections, liver regions were embedded in OCT compound (ThermoFisher, Cat# 23-730-571) in cryomolds (Sakura, Cat# 4565), snap-frozen in 2-methylbutane (Sigma-Aldrich, Cat# M32631), cooled on dry ice, and stored at −80°C.

### Mouse models

ZNRF3/RNF43^fl/fl^ mice^24^, AXIN2CreER mice and R26-LSL-EGFP mice^58^ were described previously. ZNRF3/RNF43/CTNNB1 triple floxed mice ZNRF3/RNF43/CTNNB1^fl/fl^ were generated by crossing ZNRF3/RNF43^fl/fl^ mice with CTNNB1 floxed mice CTNNB1^fl/fl^, a kind gift from Dr. Rendl lab (Icahn School of Medicine at Mount Sinai) (Jax#022775). ZNRF3/RNF43/AXIN2CreER; R26-LSL-EGFP mice were generated by crossing ZNRF3/RNF43^fl/fl^ mice with AXIN2CreER and R26-LSL-EGFP mice. AXIN2CreER;R26-LSL-EGFP mice were generated by crossing AXIN2CreER mice with R26-LSL-EGFP mice. C57BL/6J mice (8–12 weeks old) were obtained from The Jackson Laboratory.

### Dietary regimens

Mice were fed either standard chow or specialized diets tailored to specific liver disease models. All diets were provided ad libitum. A standard chow diet was provided by the Center for Comparative Medicine and Surgery at ISMMS. Chow-fed mice served as either controls (6 months) or for MASH regression studies (2 weeks or 2 months) and were also used for lineage tracing experiments (1, 2 or 9 months). To model progressive MASH with obesity, insulin resistance, and fibrosis, we used the FAT-MASH model (also referred to as the "Western" or "Fast Food Diet" model), as described previously (Tsuchida et al., 2018; Envigo, Cat# TD.120528). This regimen combines a high-fat Western diet (20–23% fat by weight; 40–45% kcal from fat, predominantly milk fat with >60% saturated fatty acids), 1.25% dietary cholesterol, glucose/fructose-enriched drinking water, and chronic low-dose carbon tetrachloride (CCl₄; 0.02 mL/kg body weight). This combined metabolic and toxic insult drives steatosis, hepatocyte ballooning, lobular inflammation, and pericellular fibrosis, already within 6 weeks. Feeding durations ranged from 6 to 18 weeks. Additionally, to model advanced MASH and fibrosis, mice were fed a choline-deficient, L-amino acid–defined, high-fat diet (CDAHFD; Research Diets Inc., Cat# A06071309i) containing 45% kcal from fat, 0.1% methionine, and no added choline. This impairs hepatic VLDL secretion and promotes steatosis, injury, and fibrosis. In the experiments using CDAHFD, the feeding duration was 9 weeks.

### AAV8-TBG-Cre–mediated hepatocyte recombination

Hepatocyte-specific genetic recombination of ZNRF3/RNF43^fl/fl^, and ZNRF3/RNF43/CTNNB1^fl/fl^ were achieved through intravenous (i.v.) injections of 10¹ -10¹¹ viral particles of pAAV8.TBG.PI.Cre.rBG (AAV8-TBG-Cre; Addgene Cat# 107787-AAV8). Control animals received an equivalent dose of pAAV.TBG.PI.eGFP.WPRE.bGH (AAV8-TBG-EGFP; Addgene Cat# 105535-AAV8). Tail vein injections were performed under sterile conditions using a total volume of 100 µL sterile PBS. Mice were euthanized at 2 weeks and 1 month after AAV8-TBG-Cre injection.

### AAV8-TBG-Cre–mediated sparse hepatocyte lineage tracing

To perform sparse hepatocyte lineage tracing, Rosa26-LSL-EGFP, and ZNRF3/RNF43^fl/fl^;Rosa26-LSL-EGFP mice carrying a Cre-dependent reporter allele (i.e. Rosa26-LSL-EGFP) were injected intravenously with low-dose AAV8-Cre to achieve sparse labeling suitable for clonal analysis. Ultra–low doses (on the order of 10 –10 genome copies per mouse, diluted in 100 µL PBS) were used to ensure clonal resolution. Mice were euthanized at 2 months and 9 months after AAV8-TBG-Cre injection for quantification of clone number, size, and zonal distribution.

### Tamoxifen-inducible AXIN2-CreER sparse hepatocyte lineage tracing

To perform sparse hepatocyte lineage tracing in AXIN2CreERT2; ZNRF3/RNF43^fl/fl^; Rosa26-LSL-EGFP mice were treated intraperitoneally with one single low dose tamoxifen injection (0.5 mg for males, 0.25 mg for females), and mice were euthanized 9 months later for quantification of clone number, size, and zonal distribution. Differential dosing was chosen to achieve comparable labeling efficiency between sexes.

### Immunohistochemistry (IHC)

Paraffin sections (3 µm) were deparaffinized in xylene (ThermoFisher, Cat# HC7001GAL) and rehydrated through graded ethanol (ThermoFisher, Cat# 22032601) followed by distilled water. Antigen retrieval was performed in 10 mM citrate buffer (pH 6.0) at 98°C for 20 min, followed by cooling on ice for 15 min. Endogenous peroxidase activity was blocked with 0.5% H₂O₂ in methanol (Sigma-Aldrich, Cat# 179337-4L) for 20 min at room temperature. Slides were rinsed in distilled water and PBS (ThermoFisher, Cat# 10010-023) for 5 min each, and hydrophobic barriers were drawn. Sections were blocked in PBS containing 5% normal goat serum (Jackson ImmunoResearch, Cat# 005-000-121) and 0.1% Triton X-100 (Sigma-Aldrich, Cat# T8787-250ML) for 30 min at room temperature. Primary antibodies (Rabbit anti-CYP2E1 1:1000, Cell Signaling Technology, Cat# 9661S; Rabbit anti-GS, 1:5000, Abcam, Cat# ab49873) were diluted in blocking buffer and incubated overnight at 4°C. After washing in PBS, slides were incubated with Histofine Simple Stain MAX PO Anti-rabbit (Nichirei Biosciences Inc., Cat# 414341F) secondary reagents for 30 min at room temperature. DAB development was performed using the ImmPACT DAB substrate kit (Vector Laboratories, Cat# SK-4105). Slides were rinsed in water, counterstained with Hematoxylin Gill III (Sigma-Aldrich, Cat# 65067-75), rinsed in tap water, dehydrated through graded ethanol, cleared in xylene, and mounted with Permount (ThermoFisher, Cat# SP15-100).

### Immunofluorescence (IF)

Paraffin sections (3 µm) were baked at 60°C for 60 min, deparaffinized in xylene, and rehydrated through 100%, 95%, 85%, and 70% ethanol, followed by distilled water. Antigen retrieval was as above (10 mM citrate, pH 6.0, 98°C, 20 min; cool 15 min on ice). After rinsing in water and PBS, hydrophobic barriers were drawn, and sections were blocked for 45 min at room temperature in PBS with 3% donkey serum (Sigma-Aldrich, Cat# S30-100ML) and 0.1% Triton X-100. Primary antibodies were diluted in blocking buffer and incubated overnight at 4°C. The following primary antibodies were used: Rabbit anti-CYP2E1 1:250 (Sigma-Aldrich, Cat# HPA009128); Rabbit anti-GS 1:2000 (Abcam, Cat# ab49873); Rat anti-Ki-67 1:1000 (eBioscience, Cat# 14-5698-82); Rat anti-CK19 1:100 (DSHB, Cat# TROMA-III; RRID:

AB_2133570); Goat anti-GFP 1:500 (Abcam, Cat# ab6673; RRID: AB_305643). Slides were washed in PBST (PBS + 0.05% Tween-20; Sigma-Aldrich, Cat# P9416) 3 × 5 min, then incubated for 30 min at room temperature with Alexa Fluor–conjugated secondary antibodies (Jackson ImmunoResearch; catalog numbers in Key Resources Table) diluted 1:500 in blocking buffer, together with DAPI (Sigma-Aldrich, Cat# D9542; 1:2000). After 3 × 5 min PBST washes and a final distilled water rinse, slides were mounted with FluoroSave (Millipore, Cat# 345789) and stored protected from light until imaging.

### Image acquisition

Brightfield, and immunofluorescence images were acquired on a Stand Axio Observer 7 Microscope (Carl Zeiss AG, 431007-9904-000) equipped with a motorized stage, high-resolution CMOS camera, and LED fluorescence illumination. Imaging was performed using 10× and 20× objectives. Brightfield images were acquired under standardized exposure and white balance conditions. For fluorescence, filter sets optimized for Cy2, Cy3, and Cy5 were used. Exposure time, gain, and illumination were optimized for each fluorophore to avoid saturation and were kept constant within each experimental batch. Images were exported as .czi, TIFF, or JPEG files for downstream analysis. Alternatively, whole slides were scanned using Hamamatsu scanner Nano Zoomer S60 combined with software NDP Scan 3.4.2 followed by analysis using Fiji ImageJ.

### Quantification of immunofluorescence (IF) and immunohistochemistry (IHC)

Quantitative analysis of immunofluorescence (IF) and immunohistochemistry (IHC) for EGFP, Ki67 and CYP2E1 were performed using Fiji/ImageJ (RRID: SCR_003070). Images were acquired under standardized magnification and illumination settings and exported as RGB TIFF or JPEG files. For colorimetric IHC stains, regions of interest (ROIs) encompassing the total hepatocyte area were selected at 4× magnification for CYP2E1 IHC quantification. The percentage of CYP2E1⁺ hepatocyte area was calculated by manually drawing positive-staining ROIs and dividing the positive area by the total hepatocyte area. Images were randomly selected to ensure representation across whole-liver sections. For IF-based analyses, Ki67⁺ hepatocytes were quantified as the number of Ki67⁺ hepatocytes per high-power field (HPF), determined by manual counting with Ki67⁺ signals cross-validated against DAPI staining. For EGFP⁺ hepatocyte clonal tracing experiments, the distribution of EGFP⁺ hepatocytes (%) in periportal (PV), parenchyma (PA), and pericentral (CV) zones was quantified as the percentage of EGFP⁺ hepatocytes per zone. PV, PA, and CV zones were defined by manually dividing the portal vein–to–central vein axis into three equal regions, as described previously^21^. GFP⁺ clones and total clone numbers were quantified by manual counting based on EGFP signal and DAPI. For batch processing, image series were imported as stacks using File > Import > Image Sequence, and .czi files were converted to JPEG using File > Save As > JPEG prior to analysis. Portal and central ROIs were defined anatomically, with periportal ROIs drawn around portal triads (PV) and pericentral ROIs drawn around central veins (CV). All quantification was performed under blinded conditions whenever feasible

### Spatial Transcriptomics

Spatial transcriptomic data were normalized using the sctransform function in Seurat, followed by dimensionality reduction with PCA (top 30 components). Public datasets and the datasets generated in this study were processed independently to avoid cross-dataset interference during normalization and feature selection. To correct for batch effects, principal components from each dataset were integrated using the Harmony package. The harmonized PCs were then embedded using UMAP (min.dist = 0.3; dims = 1:30) and clustered with the Louvain algorithm. To establish the PP–PV zonation axis, we applied monocle3 pseudo-time analysis to each sample using the harmonized PCs as input. The resulting pseudo-time values were used as an approximation of the spatial transition from periportal (PP) to pericentral (PV) regions. Differentially expressed genes between PP and PV zones were identified using the Wilcoxon rank-sum test. To assess zonation changes across conditions, we calculated the log2 fold change between PP and PV zones for each condition. The standard deviation of log2 fold change across all genes was used as a threshold to select genes showing notable variation.

These genes were subsequently categorized into three groups—less zoned, more zoned, and swapped—based on the direction and magnitude of their zonation shifts.

### DNA and RNA extraction

Genomic DNA and total RNA from tumor tissue and adjacent liver parenchyma were extracted using DNeasy Blook & Tissue Kit (QIAGEN, 69504) and RNeasy Plus Mini Kit (QIAGEN, 74134) respectively following the manufacturer’s instructions. Prior to extraction, biopsies were lysis by a TissueLyser LT (QIAGEN, 85600). Extracted genomic DNA was quantified using Qubit Fluorometer (Invitrogen). RNA concentration was measured using a NanoDrop 2000 spectrophotometer (Thermo Fisher Scientific), and RNA quality and integrity were assessed with an Agilent 2100 BioAnalyzer using the RNA 6000 Nano Kit (Agilent Technologies).

### Whole-exome sequencing (WES)

Whole-exome sequencing was performed on seven ZNRF3/RNF43 deletion tumor samples and one germline control. Whole-exome capture was performed using the V2 library (S) kit (IDT) according to the manufacturer’s guidelines. Sequencing was done on the DNBseq platform (MGI Tech). Sequence reads were aligned to the reference mouse genome GRCm38 using BWA (v0.7.17) (http://arxiv.org/abs/1303.3997). Local realignment, duplicate removal, and base-quality recalibration were performed using the Genome Analysis Toolkit (GATK, v3.6)^59^ and Picard (v2.20.0; http://broadinstitute.github.io/picard/). Median sequencing depth was 77.0× (range 73.7–84.6×). Somatic single-nucleotide variants (SNVs) and indels were called using MuTect2 (GATK v4.2.4.1)^59^ and Strelka (v2.9.10)^60^, respectively. We removed variants outside target regions; those supported by <3 Reads in the tumor; those covered by <10 reads in the tumor or <5 reads in the germline; and variants where tumor variant allele frequency (VAF) was less than 5× that of the matched germline VAF. Additional artifacts were filtered using MuTect2’s artifact detection mode in GATK 3.6. Driver genes were compiled from HCC-focused gene lists: Martincorena et al.^61^, TCGA^49^, IntoGen^62^, Schulze et al.^63^, Fujimoto et al.^64^, and the Cancer Gene Census v88 Tier 1^65^. Oncoplots were generated using the maftools R package (v2.20.0).

### RNA-sequencing data processing and analysis

RNA-seq library preparation was performed with NEBNext Ultra II directional Library Prep Kit (E7760) and sequenced on the NovaSep instrument. Sequence reads were aligned to the mouse reference genome GRCm38 using STAR (v2.7.3a)^66^ in two-pass mode. The median number of aligned reads per sample was 31.8 million (range 22.6–39.8 million). Gene quantification was performed with RSEM (v1.3.2)^67^.

Gene-level expression was analyzed with edgeR (v4.2.2)^68^. Raw expected counts from RSEM were used as input. Normalization was performed using the TMM (trimmed mean of M-values) method^69^ and differential expression was assessed using the quasi-likelihood F-test with Benjamini–Hochberg correction for multiple testing. Genes with adjusted p ≤ 0.05 were considered differentially expressed. Over-representation analysis was performed using the enricher function in clusterProfiler (v4.12.6)^70^ against the H, C2:CP: KEGG_LEGACY, C2:CP: KEGG_MEDICU, C2:CP: REACTOME, and C5:GO: BP subsets of MSigDB^71,72^, obtained via the msigdbr R package (v25.1.1). Gene sets with Benjamini–Hochberg-adjusted p ≤ 0.05 were considered significantly enriched. GSVA-based gene set scores (for liver zonation^73^, respiration, and ZRdKO/CTNNB1/MYC signatures) were computed on TPM data using GSVA (v1.52.3)^74^. Data visualization was performed using ggplot2 (v3.5.2), ggpubr (v0.6.0), and ComplexHeatmap (v2.20.0).

### Analysis of TCGA and MERiC cohorts

TCGA HCC data (clinicopathological, mutation, RNA-seq TPM) were obtained via the Genomics Data Commons^49^. Edmondson’s grades were adapted from our previous work^75^. MERiC clinicopathological, mutation, and RNA-seq FPKM data were obtained from Ng et al.^50^. GSVA scores (for liver zonation^73^, respiration, ZRdKO/CTNNB1/MYC signatures, and HCC subtype signatures^76^) were computed on TPM or FPKM data using GSVA (v1.52.3). ZRdKO-like HCCs were defined as those fulfilling both of the following criteria: (1) ZRdKO signature score in the top quartile, and (2) difference between ZRdKO score and the maximum of CTNNB1 and MYC signature scores in the top quartile (i.e., the largest ZRdKO minus CTNNB1/MYC contrast). Data visualization was performed using ggplot2 (v3.5.2), ggpubr (v0.6.0), and ComplexHeatmap (v2.20.0).

### Statistics and reproducibility

Unless otherwise stated, data are presented as mean ± standard deviation (s.d.), with n indicating the number of biological replicates (animals or independent samples). Statistical analyses for tissue-level and imaging data were performed using GraphPad Prism v10 (GraphPad Software; RRID: SCR_002798). No formal statistical method was used to predetermine sample size. Experimental groups were randomized and, where feasible, investigators were blinded to group allocation during image acquisition and quantification. Data were assumed to follow approximately normal distributions, for comparisons between two groups, two-tailed unpaired Student’s t-tests were applied (paired t-tests were used where appropriate). For comparisons among more than two groups, one-way or two-way ANOVA was performed, followed by Sidak’s or Tukey’s multiple-comparisons tests, as indicated. For non-parametric comparisons of numerical variables (e.g., signature scores between ZRdKO-like and Other HCC groups), Mann–Whitney U (Wilcoxon rank-sum) tests were used. Comparisons of ordinal variables (e.g., Edmondson grade) were performed using Mann–Whitney U tests. Comparisons of categorical variables (e.g., mutation status, Hoshida class) were evaluated by Fisher’s exact test or chi-squared test, as appropriate. Pearson correlation coefficients were used to assess relationships between continuous variables (e.g., ZRdKO signature in human

HCC, and zonation scores). RNA-seq, WES, and cohort-level statistical analyses were performed in R v4.4.1. Multiple testing correction was performed using the Benjamini–Hochberg method, unless otherwise stated. Exact test types and p-values are reported in the figure legends. When exact p-values are not listed, significance is encoded as: p < 0.05 (*), p < 0.01 (**), p < 0.001 (***), p < 0.0001 (****); ns, not significant. All tests were two-sided.

---

**Suppl Figure 1.**
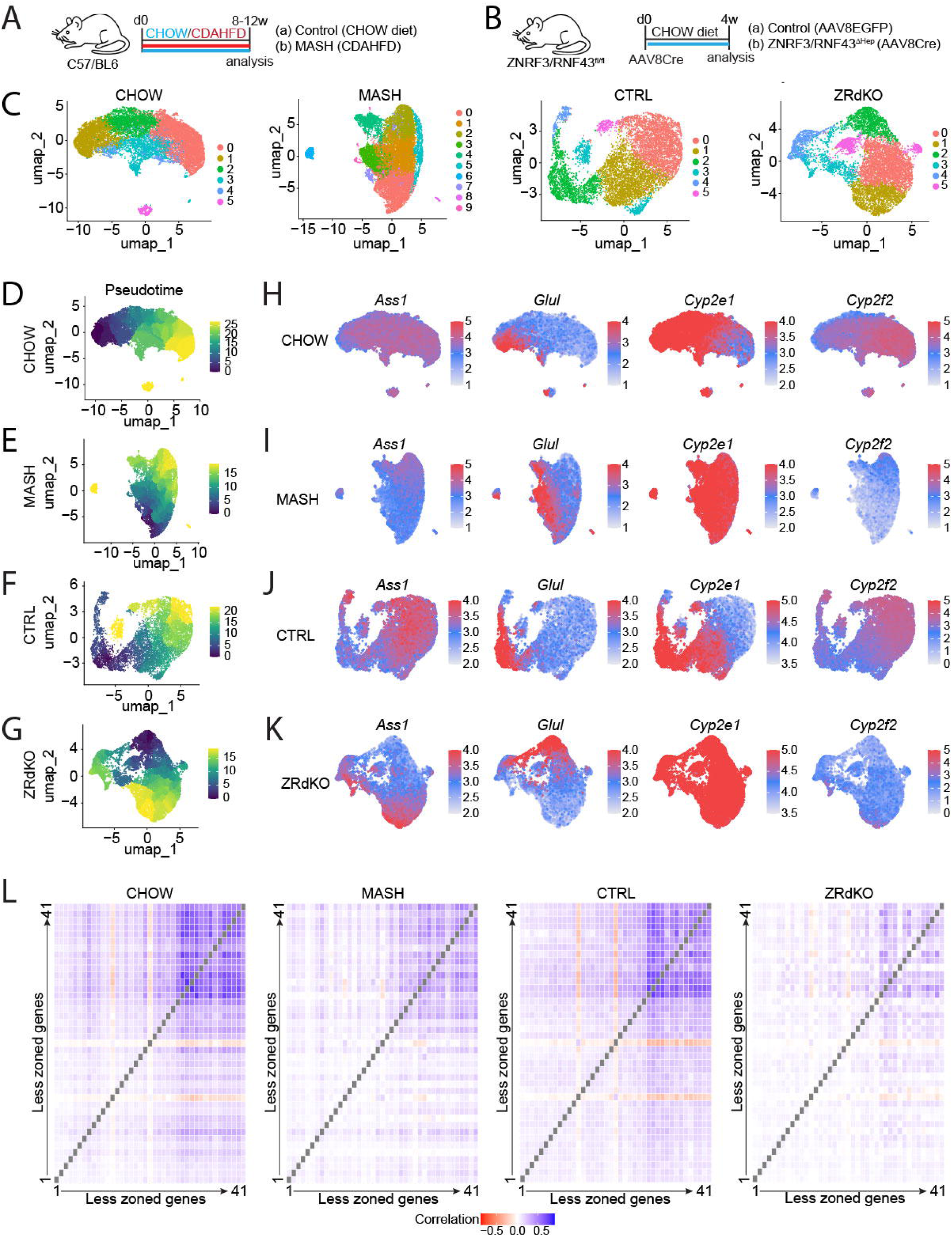

**Suppl Figure 2.**
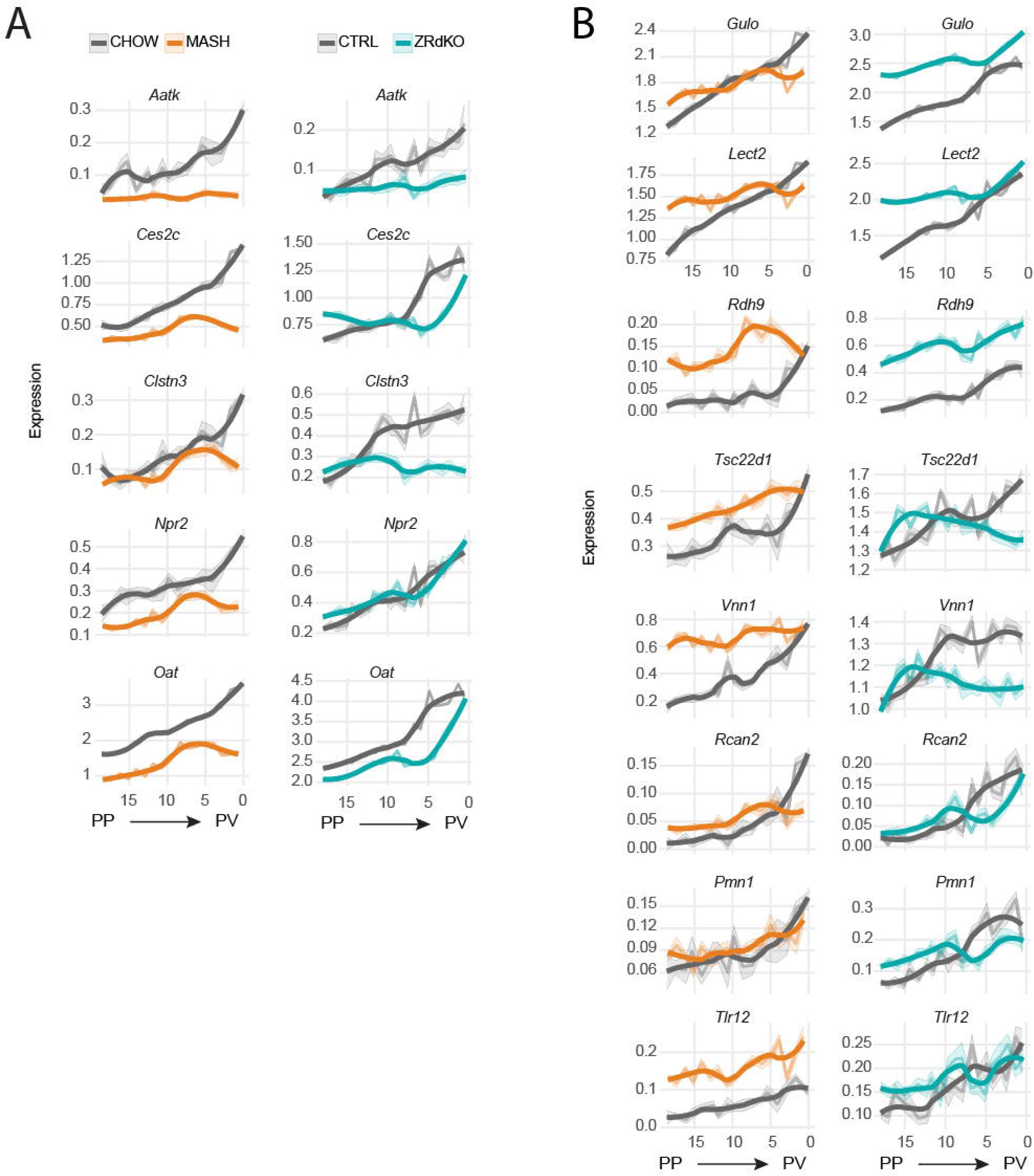

**Suppl Figure 3.**
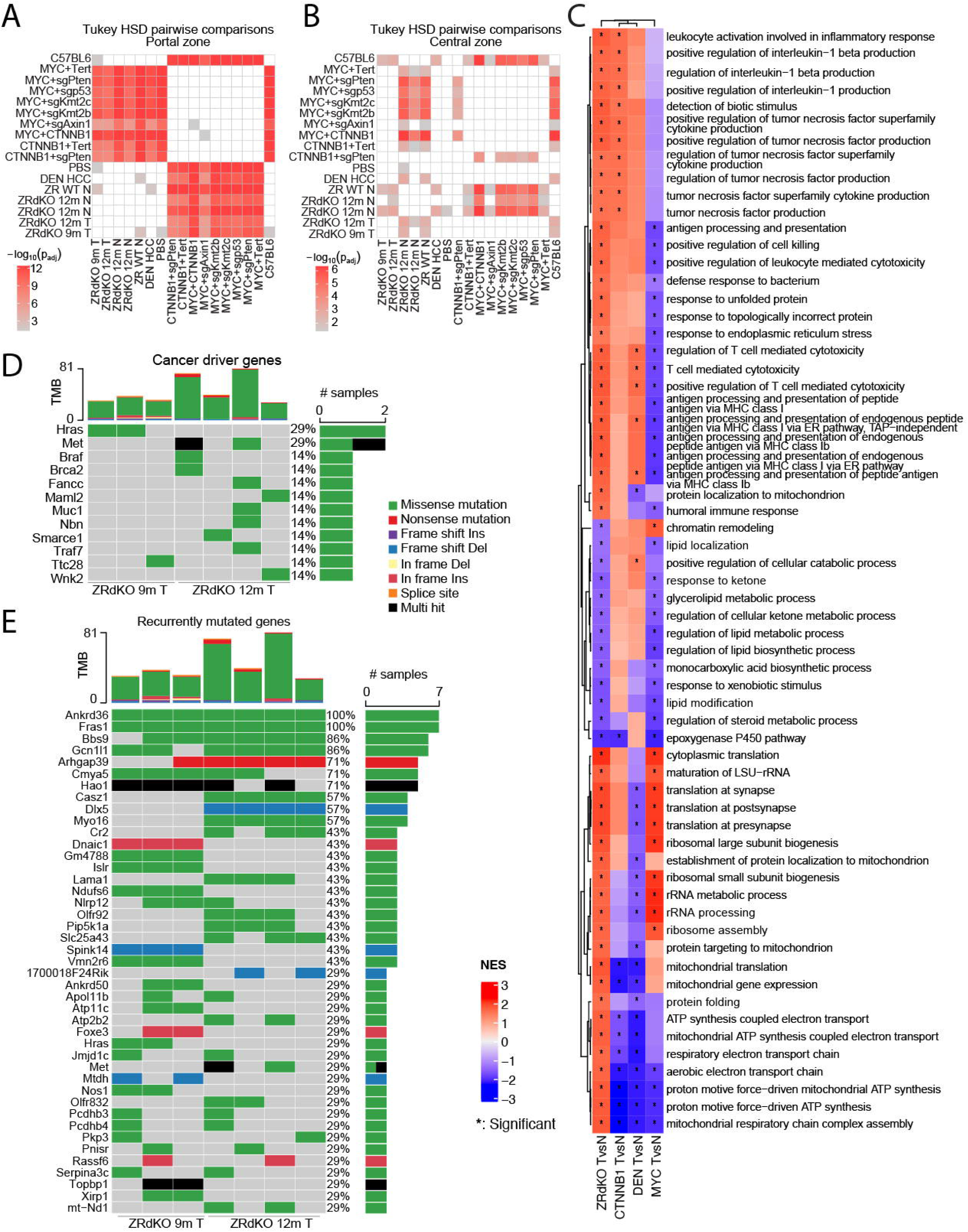

**Suppl Figure 4.**
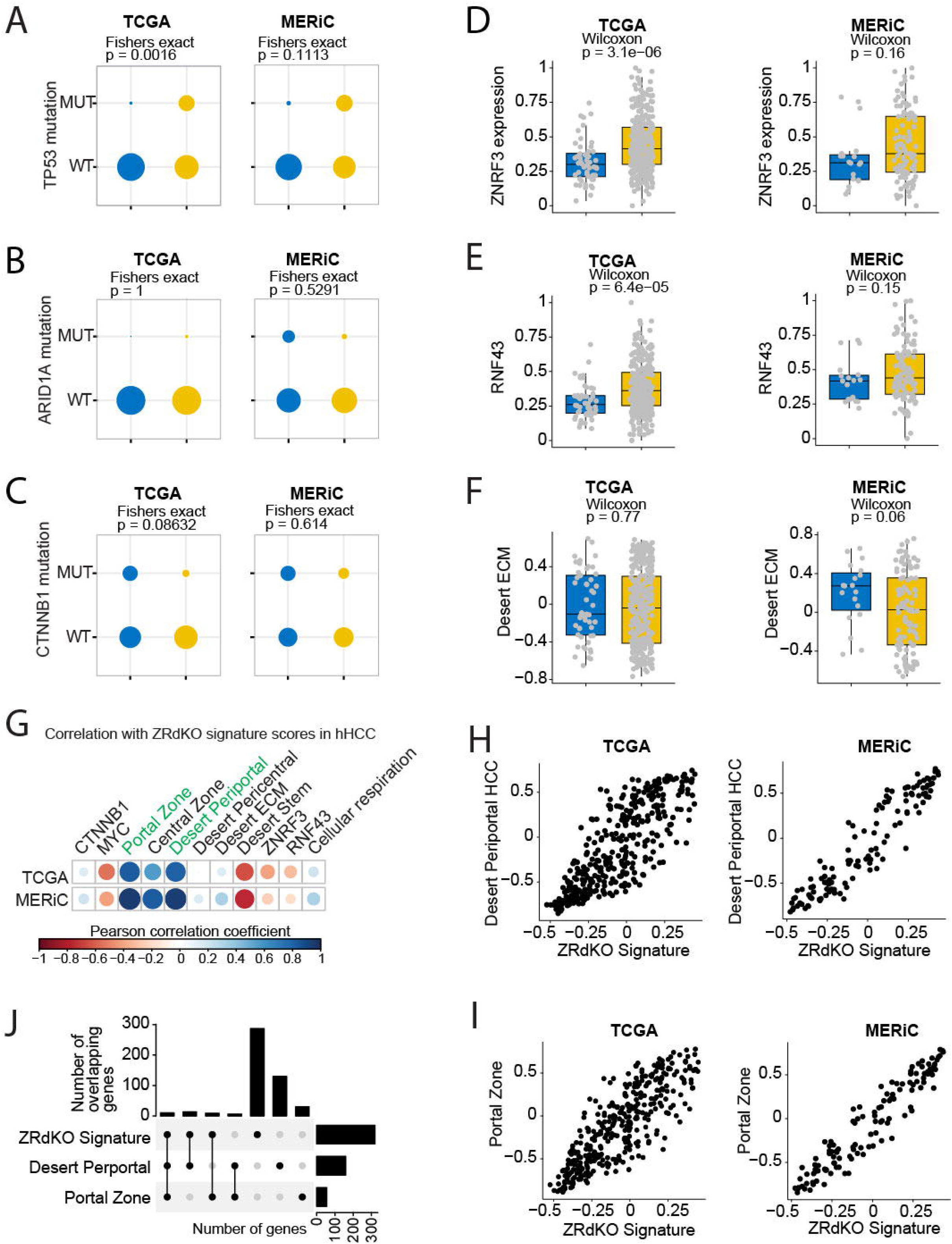

