## Supplementary material for "Liver Zonation Disruption Fuels Hepatocellular Carcinoma in Chronic Liver Disease": Suppl Legends

### Supplemental information

**Supplementary Figure 1. Spatial transcriptomics identifies shared hepatocyte zonation patterns in dietary and ZRdKO genetic liver injury model. (A)** Schematic of the dietary intervention for spatial transcriptomic profiling. C57BL/6 mice were fed either CHOW or CDAHFD for 8–12 weeks prior to tissue collection and analysis. **(B)** Schematic of the hepatocyte-specific KO experiment. ZNRF3/RNF43$^{fl/fl}$ mice were injected with AAV8-EGFP (Control) or AAV8-Cre (ZRdKO) 4 weeks before analysis. **(C)** UMAP projections of single-cell transcriptomes from CHOW, MASH, CTRL and ZRdKO samples, colored by clusters (numeric labels). **(D–G)** Pseudotemporal ordering of the same UMAPs shown in (C), presented separately for CHOW (D), MASH (E), CTRL (F), and ZRdKO (G). **(H–K)** UMAP expression maps of canonical zonation markers (Ass1, Glul, Cyp2e1, Cyp2f2) for CHOW (H), MASH (I), CTRL (J), and ZRdKO (K). Color scale ranges from low (white) to intermediate (blue) to high (red) expression. **(L)** Correlation matrices of the 41 less-zonated genes across CHOW, MASH, CTRL, and ZRdKO samples. Color scale indicates Pearson correlation coefficients from negative (red) to positive (blue).

**Supplementary Figure 2. Spatial transcriptomics identifies less zoned genes in dietary and ZRdKO genetic liver injury model.** Spatial expression profiles of representative less zoned genes along the periportal–pericentral axis expressed in arbitrary units (x axis) with decreased pericentral expression (*Aatk, Ces2c, Clstn3, Npr2, Oat*). (**A**) or increased periportal expression (*Gulo, Lect2, Rdh9, Tsc22d1, Vnn1, Rcan2, Pmn1, Tlr12*) (**B**). For each gene, expression is shown for CHOW vs MASH (left) and CTRL vs ZRdKO (right).

**Supplementary Figure 3. Expanded zonation analyses, pathway-level comparisons, and mutational features of ZNRF3/RNF43-deficient HCCs.** (**A–B**) Heatmaps of Tukey HSD pairwise comparisons of zonation GSVA scores across all samples in the ZNRF3/RNF43, DEN, and oncogenic driver studies. Panel (A) shows portal scores, and panel (B) shows central zone scores. Values represent $-\log_{10}$ adjusted *P* values. Related to Figure 5B. **(C)** Full GSEA heatmap corresponding to Fig. 5C, shown with all ontology-term labels and hierarchical clustering on both pathways and model comparisons. Asterisks denote statistically significant enrichments. (D–E) Oncoprints of somatic mutations across ZRdKO, MYC-, CTNNB1-, and DEN-driven tumors. Panel (D) shows curated cancer driver genes, and panel (E) shows recurrently mutated genes, both ordered by mutation frequency. Mutation classes are color-coded and accompanying side and top bars indicate the number of mutated samples and tumor mutational burden (total non-synonymous mutations), respectively.

**Supplementary Figure 4. Genomic, expression, and signature associations of ZRdKO-like human HCCs. (A-C)** Dot plots displaying TP53 **(A)**, ARID1A **(B)**, and CTNNB1 **(C)** mutation status in ZRdKO-like and Other HCC in TCGA (left) and MERiC (right). Fisher's exact P values are shown. **(D-F)** Box plots showing ZNRF3 expression (D), RNF43 expression (E) and ECM HCC scores (F) in ZRdKO-like and Other HCC in TCGA (left) and MERiC (right). Wilcoxon rank-sum P values are shown. **(G)** Correlation matrices showing Pearson correlations between the ZRdKO signature and other signatures shown in Figure 6L-M in TCGA (top) and MERiC (bottom). Dot color indicates correlation coefficient; dot size reflects correlation magnitude. **(H)** Shared-gene analysis for ZRdKO, periportal HCC and portal zonation signatures. Dots represent genes assigned to each signature, and connecting lines depict shared genes. Bar plots display the size of each signature and the number of overlapping genes. **(I-J)** Scatter plots showing pairwise correlations between ZRdKO signature scores and periportal-HCC scores **(I)** or portal zonation scores **(J)** for TCGA (left) and MERiC (right).
